# Label-Free In-Line Characterization of Immune Cell Culture using Quantitative Phase Imaging

**DOI:** 10.1101/2025.04.22.649403

**Authors:** Caroline E. Serafini, Viswanath Gorti, Paloma Casteleiro Costa, Aaron D. Silva Trenkle, Bharat Kanwar, Bryan Wang, Brian Wicker, Linda E. Kippner, Isaac LeCompte, Rui Qi Chen, Benjamin Joffe, Ye Li, Annie C. Bowles-Welch, Jing Li, Christine E. Brown, Gabriel A. Kwong, Stephen Balakirsky, Krishnendu Roy, Francisco E. Robles

## Abstract

Cell therapies, including T cell immunotherapies, offer promising treatments for previously untreatable diseases, but their widespread use is hindered by challenges in monitoring therapeutic cells during culture—impacting consistency, potency, and cost. This work demonstrates the use of quantitative phase imaging (QPI), specifically a compact, non-interferometric form called quantitative oblique back illumination microscopy (qOBM), for non-destructive, label-free, in-line assessment of T cell cultures. qOBM enables near real-time feedback on culture growth, contamination, and cell status (viability and activation), comparable to flow cytometry. We further apply this method to characterize genetically modified CAR T cells and explore its potential for advanced T cell phenotyping. Analysis of data from over 50 independent donors shows strong correlation between qOBM metrics and traditional destructive at-line assays. Overall, qOBM provides a powerful tool for continuous, in-line monitoring of therapeutic cell cultures, which can be transformative for improving reproducibility, reducing costs, and advancing the development of cell-based therapies.

## Introduction

Cell therapy is a promising therapeutic approach that uses living cells to treat complex diseases. These novel living therapies offer a targeted, multimodal, and personalized treatment approach that takes advantage of natural repair and surveillance mechanisms to provide long-lasting effects with significant advantages over other available therapeutic options (1, 2). Recently, T cell immunotherapies have garnered particular interest for the treatment of cardiovascular diseases, neurological disorders, and certain cancers. This has led to several US Food and Drug Administration (FDA) approved therapies and clinical trials beginning with the treatment of liquid tumors like leukemia, lymphoma, and myeloma, and more recently for the treatment of solid tumors, including tumors of the brain, liver, pancreas, lung, breast, and kidney, to name a few (3–9).

While T cell immunotherapy holds significant promise, patient access to these new treatment is limited due to high costs and variability associated with the cell culture process (10). In typical biological labs, for example, cells are cultured in a monolayer, which can be easily monitored non-invasively to ensure quality. However, cells grown in a monolayer do not produce a sufficient number of cells to be used as a therapeutic. As such, cells intended for use as a therapeutic are often grown in suspension in bioreactors, which enable scalable cell manufacturing (9, 11–13). Unfortunately, a persisting issue with these platforms is the inability to monitor cells inside the bioreactor without killing or removing the cells from their sterile environment (i.e., in-line monitoring), making it difficult to monitor cells and ensure their quality throughout the culture process.

The gold standard for cell characterization and culture monitoring is fluorescence-based technologies like flow cytometry, immunohistochemistry, and immunofluorescence. All three of these modalities utilize fluorescent antibodies which label individual cells based on their surface protein expression. As illustrated in Figure 1(A), the cells inside bioreactors must be removed from the culture and prepared for flow cytometry which may include re-suspending cultures to a particular density, additional incubation time, and/or cell fixation. The cells are then labeled with antibodies to identify live cells, T cells (CD3^+^ antibody), activated cells (CD69^+^ or CD25^+^ antibodies for early and late stage activation, respectively), and cells with specific subtypes (CD4^+^ or CD8^+^ antibodies for helper and killer T cells, respectively), among other markers. Finally, the cells can be imaged/detected with an excitation laser and sorted to determine the population of cells exhibiting a particular marker. While these modalities do provide accurate measurements of the cell population and characterization of the bioreactor cell culture at the time of sampling, they are typically endpoint assays and require cells to be fixed and/or labeled with exogenous contrast agents (14–16). This means that cells must be removed from the culture, which can hinder cell expansion (especially when cells are removed at time points early on in the culture). Further, opening/probing the culture vessel to take a sample can introduce contaminants that can terminate the cell culture. With these limitations, monitoring cell cultures is a rather arduous task, which significantly limits the number of measurements that can be made and effectively prevents continuous monitoring and characterization of cell cultures.

**Fig. 1.**
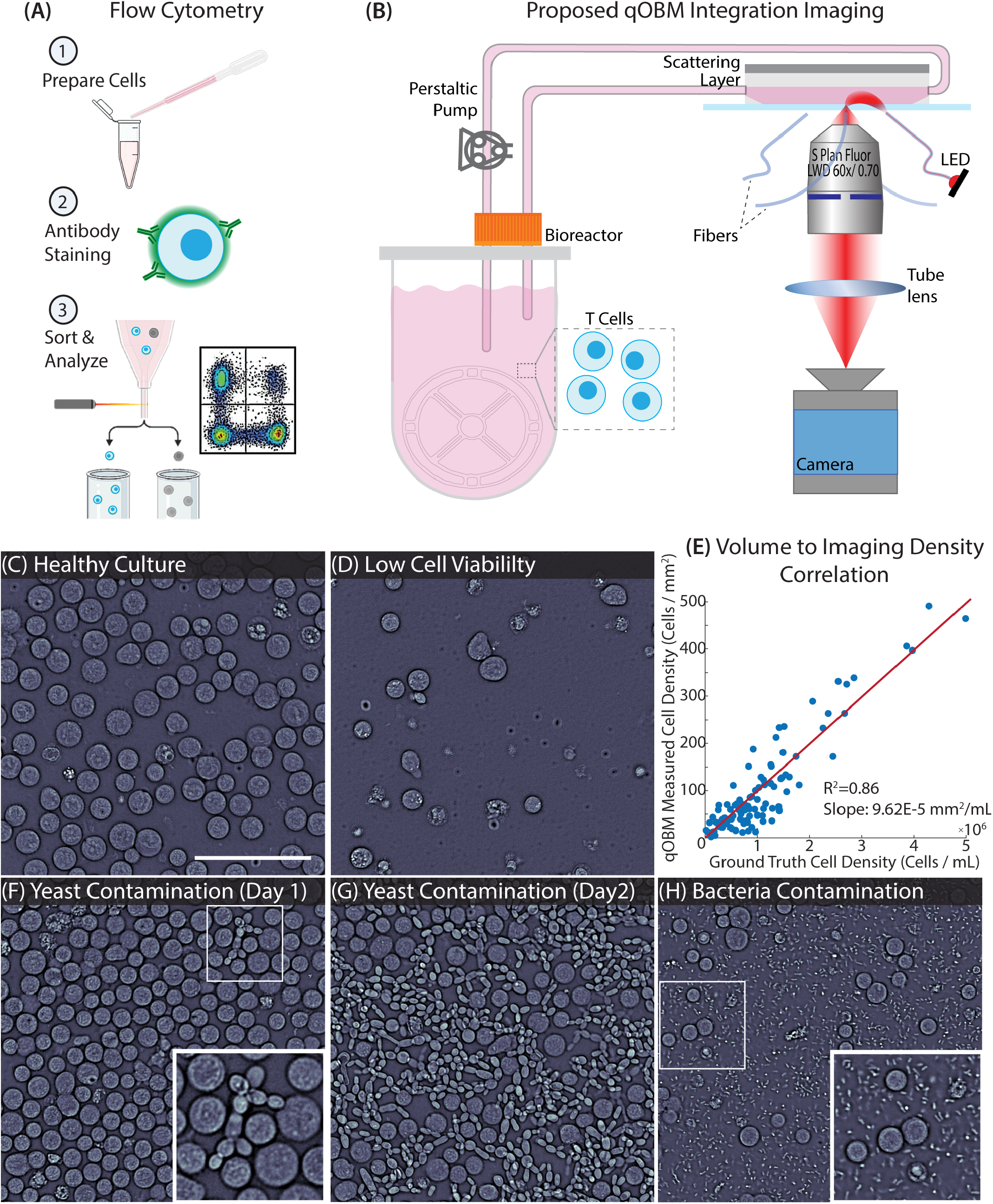
qOBM enables visualization of cells growing inside a bioreactor: (A) Flow cytometry, the gold standard for characterizing cells in bioreactors, requires (1) cell preparation, (2) surface staining with a desired antibody, and (3) sorting and analysis. (B) An overview of the qOBM integrated imaging system. This includes cells flowing out of the bioreactor through a custom flow cell. The cells are then imaged by the qOBM system illuminated sequentially by four 720 nm LEDs. The cells are then returned to the bioreactor for continued culture. (C) Healthy, growing T cell culture. (D) Low viability T cell culture. (E) Correlation between measured cell density with flow cytometry [Cells / mL] and the qOBM imaged cell density [Cells / mm^2^]. (F) T cell culture with yeast contamination on day 1 of culture and (G) 24 hours later. (H) T cell culture with bacterial contamination. Scale bar is 100 µm.

To circumvent the need for exogenous contrast agents, label-free imaging modalities like quantitative phase imaging (QPI) and autofluorescence lifetime imaging microscopy (FLIM) have been employed. Such technologies can characterize critical cell culture attributes, including cell viability (17) and subtypes (CD60^+^, CD4^+^, CD8^+^) (18). However, to date, such efforts have required cell samples to be removed from the cell culture vessels for analysis, primarily because these methods often require bulky systems with relatively expensive and complicated optical equipment, which are not particularly well suited for in-line monitoring of cell therapeutics. Thus, there remains a significant unmet need for non-destructive, label-free, in-line analysis methods to monitor and characterize cultured immunotherapy T cells.

In this paper, we introduce quantitative Oblique Back-illumination Microscopy (qOBM) to enable label-free, noninvasive, continuous in-line monitoring of T cell immunotherapy cultures grown in bioreactors. qOBM achieves quantitative phase imaging using principles of oblique, partially coherent illumination, (19–22) which does not require a dedicated interferometer to recover phase information (and refractive index properties), unlike traditional QPI technologies. Indeed, qOBM was originally developed to image thick samples,(21–25) but this alternative approach to QPI also provides advantages for imaging thin samples as the system is simple (no interferometer), compact, and low-cost, making it well-suited for in-line monitoring of cell cultures. As illustrated in Figure 1(B), our approach comprises flowing T cells out of the bioreactor into an imaging flow cell where qOBM captures quantitative phase information of the cell culture. The cells are then returned to the culture vessel for continued expansion. The system is compact and can fit inside typical incubators. By addressing a critical unmet need in cell manufacturing, qOBM offers the ability to image and characterize T cells in-line during the manufacturing process while maintaining culture sterility. The results of this work demonstrate that qOBM offers an in-line assay capable of determining T cell culture viability and activated T cells. We further apply this work for the more clinically relevant characterization of genetically modified chimeric-antigen receptor (CAR) T cells, and the possibility of CD4^+^ and CD8^+^ phenotyping is discussed.

## Results

### A. qOBM imaging provides real-time insight into T cell culture

A compact qOBM imaging system was developed and integrated within a cell incubator for imaging cells inside a bioreactor. The system measures 8in x 12in x 10in, which can fit into nearly any cell incubator with an access port (see methods section). For imaging, cells are transported from a vertical wheel bioreactor into an imaging flow cell with a scattering layer on top (see methods section) via closed-loop tubing and a peristaltic pump. After imaging within the flow cell (acquisition rate of ∼ 10Hz), cells are flushed back into the bioreactor with cell culture media for continued expansion. The compact, epi-mode nature of qOBM makes it an ideal QPI system for integration inside of the incubator due to the small footprint of the microscope. Further, qOBM utilizes low-cost LEDs coupled to multimode fibers for illumination. The LEDs (and all electronic controls) are located outside of the cell incubator (thus, they are immune to the effects of humidity). This allows for simple illumination configuration without the need for lasers, or more complex illumination boards that could also be implemented for oblique, partially coherent illumination (20). Finally, the low-cost nature of the qOBM system makes it a widely adoptable system for in-line monitoring of cell culture. In fact, any brightfield microscope (a commonplace tool in cell culture labs) can be retrofitted to a qOBM system for <$500 (see Supplemental Table S1); (26) thus this technology is widely adaptable for most labs to use for cell culture monitoring.

As seen in Figure 1(C-F), qOBM provides high-resolution images of cells during culture. These images can be acquired continuously and in real-time; thus, qOBM can provide non-destructive feedback at any regular interval during the cell culture process. Figure 1(C) shows a high-viability cell culture where cells are proliferating and expanding at a high rate, characterized by the relatively high density of cells in the image. This is in contrast to Figure 1(D) where cells are growing at a lower rate with lower viability, which is characterized by the lower density. We characterize this relationship in Figure 1(E) where we plot the ground truth cell density in cells / mL from flow cytometry versus the density of qOBM imaged cells in cells / mm^2^. With this, we quantify the relationship between qOBM imaging density and cell density to be the slope of the line: 0.0001 mm^2^/mL and see an R^2^ value of 0.86 when imaging a minimum area of 0.25 mm^2^. (Imaging more areas would reduce the error – more on this below.) Being able to visualize and quantify cell density is useful, as adjustments to the culture input parameters (e.g., glucose, lactate) can be made in real-time to ensure high yield and high quality of the cell therapeutics.

Finally, qOBM can also provide insight into contaminants. Figure 1(F), for example, shows a few yeast cells that have contaminated the culture. The progression of this contamination can be seen in Figure 1(G) which was taken 24 hours later when the yeast contaminant had clearly infected the entire culture. Interestingly, qOBM was able to detect the contamination 12 hours before other common in-line sensors (e.g., oxygen) detected anomalous conditions. Figure 1(H) shows another example of contamination, in this case, bacteria. These contaminants are smaller and can be more challenging to detect and differentiate from debris, but qOBM is able to resolve these organisms with high contrast. Supplemental Video SV1 also shows how qOBM can visualize the bacteria moving around the sample. These bacteria closely resemble prior qOBM research visualizing microbes (27).

These results clearly show that qOBM can be used to visually monitor cell cultures non-destructively in-line. In the following sections, we show how these imaging data can be processed to quantify important cell culture parameters.

### B. A qOBM imaging assay quantifies cell culture viability

Here we develop a qOBM phase imaging assay to quantify cell culture viability, and we compare the accuracy of the results to conventional end-point flow cytometry (taken as ground truth).

First, we sought to determine how individual dead cells could be identified with qOBM. With this first goal in mind, we modified the qOBM phase imaging system to enable simultaneous detection of a fluorescence-based live/dead imaging assay. This was achieved by adding a 385 nm illumination channel to the qOBM system to serve as the excitation source for DAPI (4’,6-diamidino-2-phenylindole), enabling near simultaneous multi-modal imaging (phase and fluorescence) with perfectly co-registered images of the two modalities. DAPI is a fluorescent dye that binds to DNA that is typically used to stain the nuclei of fixed cells. However, in live cells and at low concentrations (1µg/mL), DAPI is not able to permeate through the membrane of live cells, and can thus be used as a live/dead assay (28, 29). As seen in Figure 2(A), the quantitative phase qOBM image (left) and the DAPI-fluorescence image (right) can be overlaid to gain ground truth regarding live and dead cells.

**Fig. 2.**
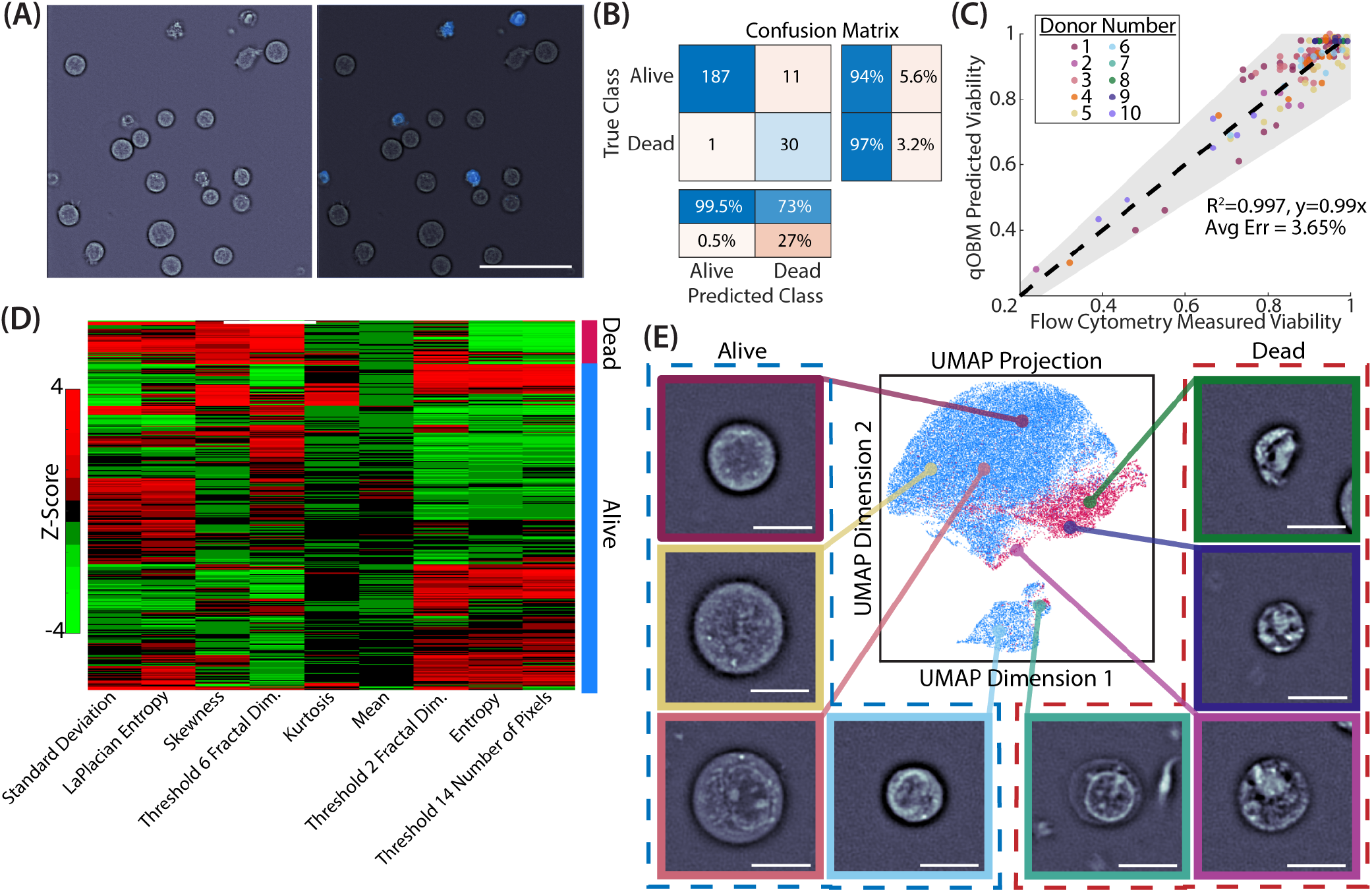
Quantification of culture viability: (A) Example cells stained with DAPI fluorescence. Left image shows the qOBM phase image and the right image shows the fluorescence overlaid with the phase. Blue cells are dead cells stained with DAPI. Scale bar is 50 µm. (B) Confusion matrix of a live/dead image-based assay trained using DAPI stained cells. (C) Network results of the qOBM predicted culture viability vs flow cytometry tested on cultures from 10 unique donors. (D) Clustergram demonstrating the correlation of curated image features from live and dead cells. (E) A UMAP of all tested live and dead cells where distinct clusters of live and dead cells form. Representative images of the cells within the clusters are also shown. This UMAP was created based on the features seen in (D). Scale bars are 10 µm.

With the ground truth live/dead fluorescent labels, next we sought to create a neural network to identify the live/dead cells using only the phase images which can be applied inline without labels. For this task we applied a residual neural network (ResNet), which can train quickly on complex imaging data and can evaluate intricate details included in the images for classification (30). Our network consisted of 30 epochs and 104 layers. The network was given 763 cells (103 dead, 660 alive) in 256×256 pixel input images of the phase values of masked cells. The network created using a 70:30 training:test split. The testing of that ResNet (as seen in the confusion matrix in Figure 2B) demonstrated a 94.7% accuracy.

Finally, we applied the trained network across cultures from 10 different donors (with images acquired at-line without labels) from which we had ground truth viability percentages from flow cytometry. These data comprised a total of 137 independent culture measurements (from the 10 individuals) each with >100 cells for a total of 38,917 cells classified as alive or dead by the network. As seen in Figure 2(C), this testing set revealed an average error of 3.94% of the qOBM determined viability versus that of the flow cytometry. Further, the datasets exemplified near-perfect correlation with an R^2^ value of 0.997. We emphasize that this analysis works well across 137 culture measurements from 10 unique donors (both from commercially available cell lines and cells from peripheral blood draws). Supplemental Tables S2-S4 summarize the donor sources and number of cells used.

While the network performed exceptionally well, we wanted to further investigate what types of features of the cells were used to determine whether a cell was alive or dead. As seen in Figure 2(D), as well as the representative images of live and dead cells in Figure 2(E), we can see that dead cells tend to have greater variability in their refractive index (RI) distribution with higher RI-valued clusters compared to their live counterparts. These are reflected in the features of phase standard deviation, skewness, kurtosis, entropy, and the fractal dimension features (see Methods section for details regarding feature calculation and selection). The features in Figure 2(D) were then used to create the UMAP as seen in Figure 2(E). Here, we see the live and dead cells clearly separate. We also see unique clusters in each of these populations including larger versus smaller live cells and dead cells with varying levels of nuclear contrast, size, and shapes. These different features may be due to a variety of conditions including culture activation, how long the cells have been dead, the method of cellular death (i.e. apoptosis versus necrosis), trauma to the cell during the cell circulation or harvesting process, etc. While these features are different on a cell-by-cell basis, as seen in Supplemental Figure S1, we can see the homogeneity that exists among an entire culture of cells; thus, we can apply this model with confidence on cells from different donors.

### C. A qOBM imaging assay quantifies cell culture activation

After determining culture viability, we sought to utilize the qOBM images to further characterize the activation state of cell cultures. In a culture, T cell activation mimics the response that T cells undergo in the body when exposed to antigens or pathogens. To accomplish this in vitro, specific antigens, mitogens, or monoclonal antibodies can be added to the culture to activate T cells. Upon exposure to the stimuli, T cells transform from their naive, quiescent state to take on activated immune response functions (such as CD4^+^ helper cells and CD8^+^ killer cells) (31–33). As described in detail in the Methods, the cells used in these experiments were activated with CD3/CD28 and IL-2.

We collected images biweekly from cell cultures growing from Day (D) 0 to D21 of culture. qOBM images of a representative cell culture can be seen in Figure 3(A-F). We note that the D0, naive cells in Figure 3(A) had not yet been exposed to activation beads. The cells in Figure 3(B-D) D3-D10 demonstrated high CD25^+^ measured activation (>97%) –CD25^+^ activation was used as it is a measure of sustained activation and is more relevant for D3+ cells (34). The cells in Figure 3(E) D14 showed a mixture in their activation state with 34.7% activated. Finally, the cells in Figure 3(F) D21 are post-activation cells with low CD25^+^ activation after previously undergoing activation (CD25^+^: 6.56%). Visually, we found that the morphologies of quiescent, post-activation cells are slightly different than naive cells (see Supplemental Figure S2). Despite potential differences, both groups ultimately are characterized as quiescent, CD25^-^ cells; thus, to create a network that can accurately characterize a given cell culture, the quiescent group of cells should contain both naive and post-activation cells. Further, T cell cultures manufactured as therapeutics contain some form of activating media, and thus the post-activation cells are more relevant for culture characterization than naive cells. In our time-lapsed images of the cell culture, clear visual differences can be seen over time: Activated cells in D3-10 (Fig. 3B-D) have larger sizes than naive cells in D0 (Fig. 3(A)). Activated cells have clear nuclear contrast and seemingly higher heterogeneity within the culture. By D14 (Fig. 3(E)), we see a heterogeneous mix of large, activated cells and smaller, quiescent cells. Finally, by D21 (Fig. 3(F)), we see an increased number of smaller, quiescent, post-activation cells.

**Fig. 3.**
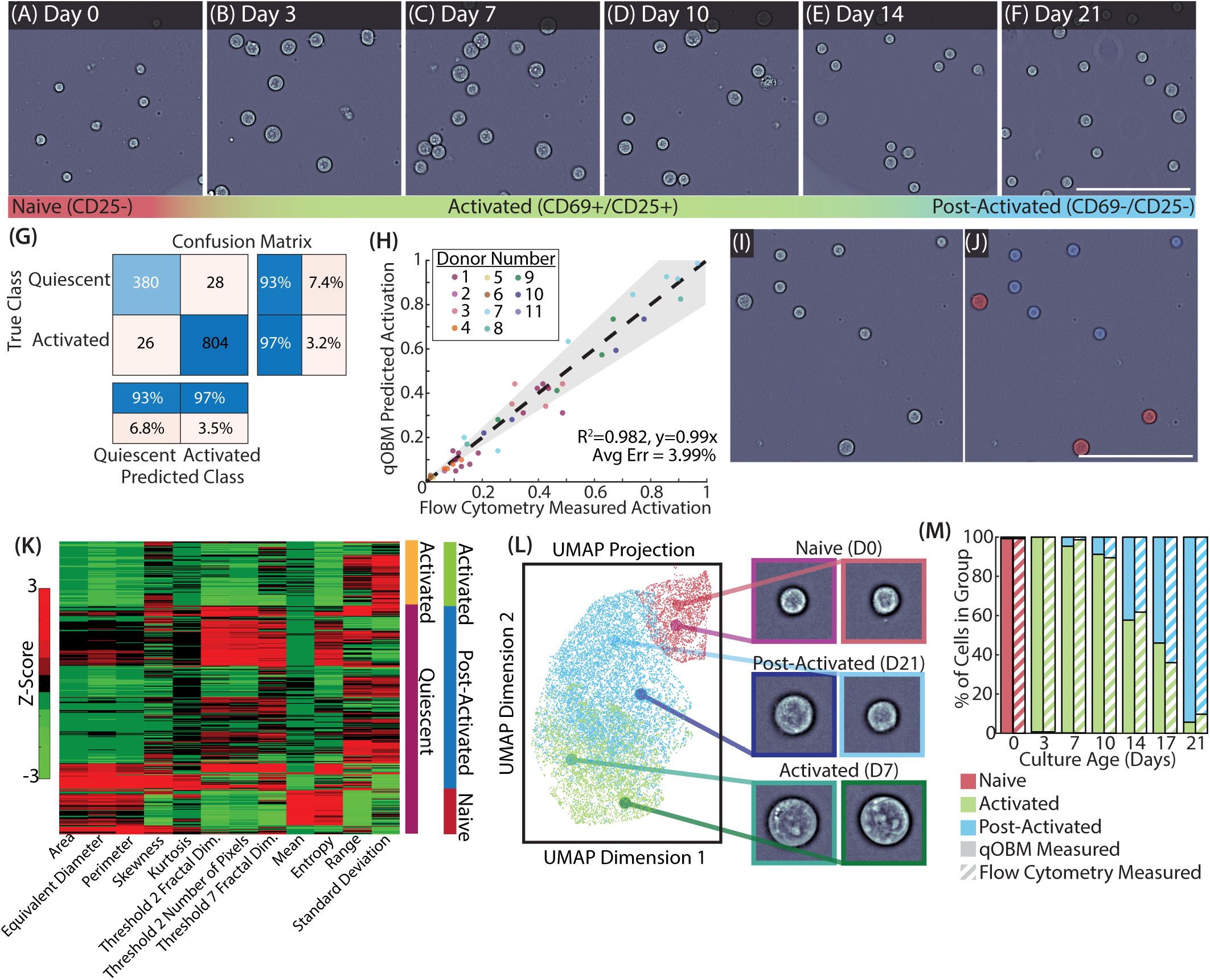
Quantification of culture activation: (A)-(F) qOBM phase images of cell cultures from naive D0 cells to activated D3-10 cell cultures, a partially activated culture on D14, and a post-activation D21 culture. (G) Confusion matrix of activated and quiescent cell cultures classification using qOBM phase images. (H) Scatter plot of the qOBM predicted culture activation vs flow cytometry measurements tested on cultures from 11 unique donors. (I-J) qOBM phase image and phase and fluorescence overlay. Blue cells are quiescent and the red cells are activated. (K) Clustergram showing the correlation of curated image features of activated and quiescent cells. (L) UMAP of all tested activated, post-activation, or naive cells which shows distinct clusters of the three groups. Representative cell images from each cluster are also included. This UMAP was created based on the features seen in (K). (M) Percent stacked barchart of the percentage of cells in each group by day of culture (solid) compared to those measured with flow cytometry (striped). Scale bars are 100 µm.

In an effort to create a network capable of classifying cells as activated or quiescent, we created a second ResNet. Using a similar pipeline as that described for the viability network, we used quiescent cell image crops from cultures with < 5% of activated cells for training. These cultures included cells never exposed to activation beads (naive cells), and D21 post-activation cells (which were no longer activated). In contrast, the activated cultures used for training had an activation of > 95%. These cultures consisted of D3-10 cells. This network was trained using a total of 956 quiescent cells and 1,936 activated cells from 3 unique donors. Results show a network accuracy (training set) of 95.4%. A confusion matrix of the outputs of this network can be seen in Figure 3(G). Supplemental Table S4 summarizes the donor source.

We then tested this network on cell cultures from 11 unique donors containing a total of 18,752 cells. While we lack a cell-to-cell ground truth regarding the activation state of the individual cell, we can compare the qOBM predicted activation percentage to that measured using flow-cytometry, which we take as ground truth (CD69^+^/CD25^+^). We note that CD69^+^ was used for D0 cells as a marker of early stage activation and CD25^+^ was used for all other cells (D3+) as a marker of late stage activation. This protocol is consistent with literature to determine the appropriate activation of T cells (35). Results again show an extremely high correlation of the qOBM predicted value compared with the ground truth CD69^+^/CD25^+^ activation, as visualized in Figure 3(H). We see an average error of 3.99% and an R^2^ value of 0.982. Qualitative evaluation of this network can be seen in Figure 3(I&J), where we see the (input) phase image in (I) and the (output) network classification image in (J) with the 3 larger, CD25^+^ cells labeled in red and the 6 smaller, CD25^-^ cells labeled in blue.

As done with the viability assay, we extracted cellular features to understand what types of structures differentiate activated and quiescent cells. The most significant features are seen in Figure 3(K), which shows that activated cells are larger, contain more heterogeneity (i.e. larger phase value standard deviation and lower entropy), and contain greater amounts of lower RI content when compared to quiescent cells. Next, the features in Figure 3(K) were used to generate the UMAP in Figure 3(L). Here, we can see unique clusters visualizing the quiescent cells on the top and the activated cells in the bottom cluster. With this, we can clearly distinguish the activation state of T cells in culture. Further, using known time that the cultures were exposure to activation media, we can label quiescent cells as naive or post-activation cells. Indeed, some variability among the quiescent cell populations can be observed, including differences in cell size and RI values. Remarkably, the UMAP also shows distinct clusters between the post-activation and naive cells, with the post-activation cells mapping between the activated and naive cells, and the post-activated quiescent cells showing more overlap with the activated cells. This behavior is expected given cells gradually transition from activated back to quiescent (post-activation).

Finally, Figure 3(M) shows the average activated and quiescent cell populations from D0 to D21 as determined by the qOBM phase-imaging assay (solid bars) and flow cytometry (striped bars), averaged across the 11 cell donors. Both measurements clearly show the same trends, starting with D0 where ∼ 100% of the T cell population is quiescent (naive), by D3 nearly all cells are activated, but then the number of quiescent, post-activation T cells increases, until D21 where quiescent, post-activation T cells make up nearly the entire cell population. Importantly, there are no statistically significant differences between the qOBM imaging-based assays and flow cytometry. For a full breakdown of the number of cells included here and the statistical analysis, see Supplemental Figure S3.

### D. qOBM viability and activation assays applied for inline characterization of T cells grown in bioreactors

With networks demonstrating high accuracy in characterizing culture viability and activation, we sought to apply these networks to characterize T cells in-line. The pipeline for classifying cells is as follows: First, cells are segmented using the Segment Anything Model (SAM) that was recently implemented to build a cell segmentation algorithm (36, 37). Next, the phase values of each cell were classified with the viability residual network as live or dead. Finally, cells characterized as live were fed to the activation residual network to classify them as activated or quiescent and estimate culture activation. This work flow can be seen visually in Figure 4(A).

**Fig. 4.**
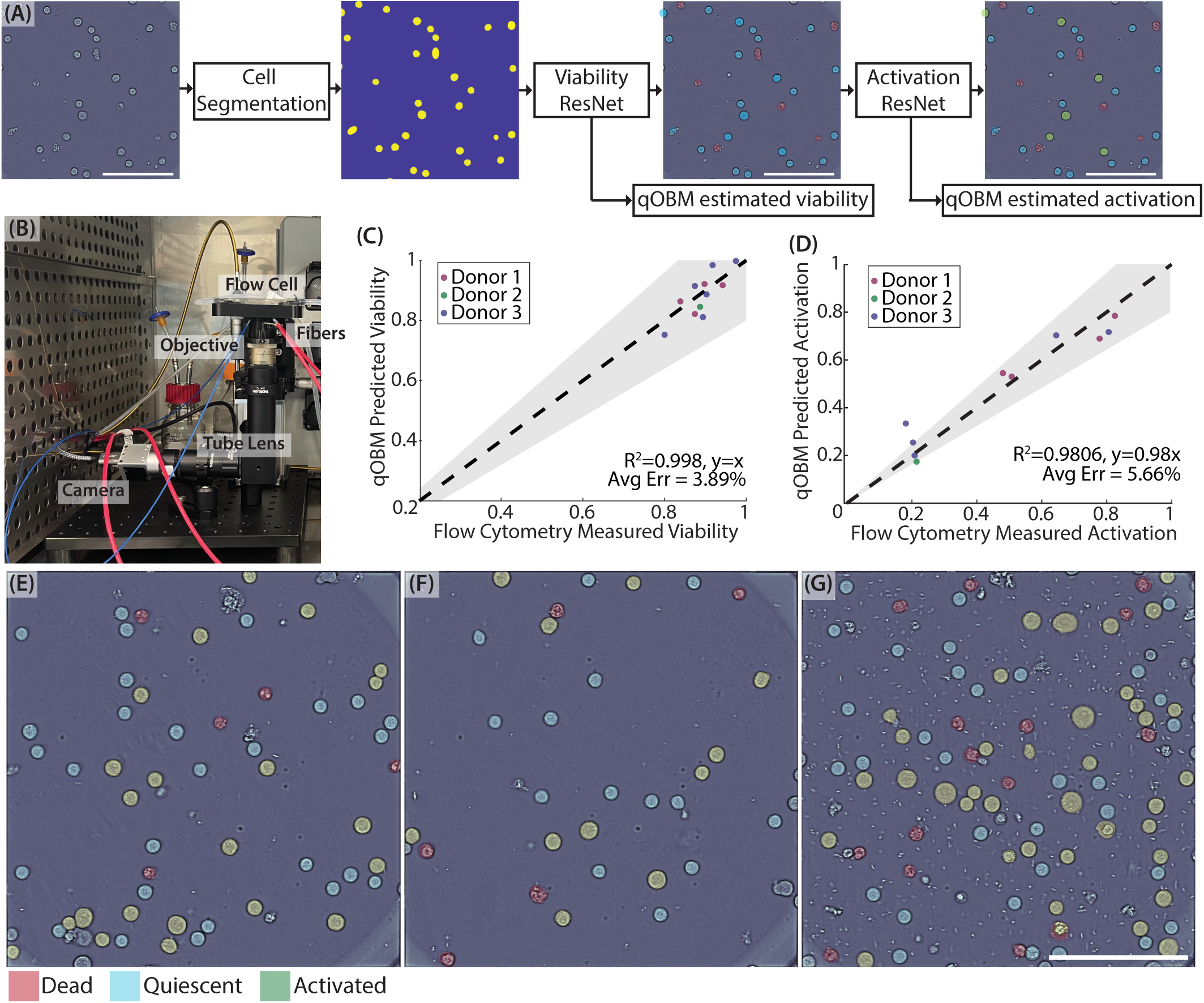
Characterization of T cell cultures in-line: (A) Flow chart depicting the process in-line T cells characterization with qOMB. First, cells are segmented, then the viability residual network is applied to determine culture viability, and finally, the activation network is applied to the live cells to determine the culture activation. (B) Photograph of the qOBM system integrated into the incubator. (C) Scatter plot of the in-line qOBM measured viability vs flow cytometry. (D) Scatter plot of the in-line qOBM measured activation vs flow cytometry. (E)-(G) Representative images of classified T cells acquired in-line. Cells highlighted in red are dead, in blue are quiescent, and in green are activated. Scale bars are 100 µm.

To collect these data, we utilized the compact, incubator compatible qOBM system, as described in the Methods H and (38). Figure 4(B) shows a photograph of the system inside the incubator with a flow cell atop the sample stage. Cells then flow into the flow cell from the bioreactor at a rate of 4 mL/min, and after ∼ 3 minutes the circulation pumps are turned off. We then allow 1-5 minutes for cells to settle to the bottom on the flow cell where they are then imaged by the compact qOBM system. One to five qOBM images are acquired to ensure each “culture measurement” uses at least 100 cells for evaluation. For each of these images, a motorized translation stage changes the field of view to image different cells without re-engaging the circulation pumps. Note that all optical equipment is located within the incubator including the microscope objective, tube lens, and camera; but all other electronics associated with the system (including computers, LED modules, and Arduino controls) are housed outside of the incubator to prevent issues with humidity. The incubator system allows cells to remain at their optimal temperature and humidity for continued growth.

We then compared the output of our qOBM predicted viability and activation to that of samples taken with flow cytometry from 10 independent culture measurements from 3 different donors. Samples are taken for flow cytometry from a sampling port in the tubing prior to qOBM imaging. The qOBM predicted viability vs the flow cytometry measured viability shows an R^2^=0.998 with an average error of 3.89% (Figure 4(C)), and the qOBM predicted activation vs the flow cytometry measured activation shows an R^2^=0.9806 with an average error of 5.66% (Figure 4(D)). These remarkable results show the potential of qOBM to accurately characterize T cell cultures in-line.

Finally, in an effort to visualize the cultures, qOBM images can be colorized to show how individual cells are classified. Figures 4(E-G) show cells classified as dead in red, quiescent cells in blue, and activated cells in green. This process is fast and can be performed in near-real-time for visual validation. A 300*µ*m x 300*µ*m field-of-view with 150 cells can be segmented, classified, and colorized in 46 seconds using a regular bench-top computer. This allows researchers to gain feedback regarding their samples quickly, easily, and accurately without the need to perturb the culture.

### E. qOBM image-based viability and activation assays applied to genetically modified CAR-T cell samples

One particular immunotherapeutic of recent interest is chimeric antigen receptor (CAR)-T cells (39–42). This innovative therapeutic strategy involves engineering a patient’s own T cells to express synthetic receptors, termed CARs, which are designed to recognize specific antigens present on tumor cells. Once added back into the patient’s body, these CAR-T cells become targeted immune cells, capable of selectively destroying cancer cells (39, 40). In an effort to demonstrate the generalizability of the qOBM images-based assays, we sought to characterize cultured CAR-T cells with the same developed networks.

Ninety culture measurements were collected using CAR-T cells from 43 separate donors (a total of 26,657 cells) (43) where the cells were cultured in vitro and analyzed. First, to demonstrate the similarities between genetically modified CAR-T cells and the untargeted donor T cells used in the analysis above, we extracted the same features from the CAR-T cells and plotted them on a UMAP with the original T cells. As seen in Figure 5(A), the live CAR-Ts in light blue plot directly on top of the untargeted live T cells in darker blue, and the dead CAR-Ts in light pink plot directly on top of the dead untargeted T cells in red. This demonstrates the similarity of the phase values and structure of the cells and the ability of the previously trained network to be applied directly to the CAR-T cells. Interestingly, we see a small cluster of cells in dark blue that map to the bottom of the UMAP, which from our previous analysis, we now understand that these cells are quiescent, unactivated T cells who have not gone through the activation process. All CAR-T cell samples were activated; and thus, we do not see any CAR-T cells that plot in that region. Further, we applied the viability network trained on the DAPI-stained T cells to the CAR-Ts. As seen in Figure 5(B), the qOBM viability assay has a high positive correlation with R^2^=0.998 and an average error of 2.27% compared to flow cytometry measurements. With these remarkable results from 43 independent donors, we demonstrate that the qOBM viability assay can be applied to the therapeutically relevant, engineered CAR-T cells.

**Fig. 5.**
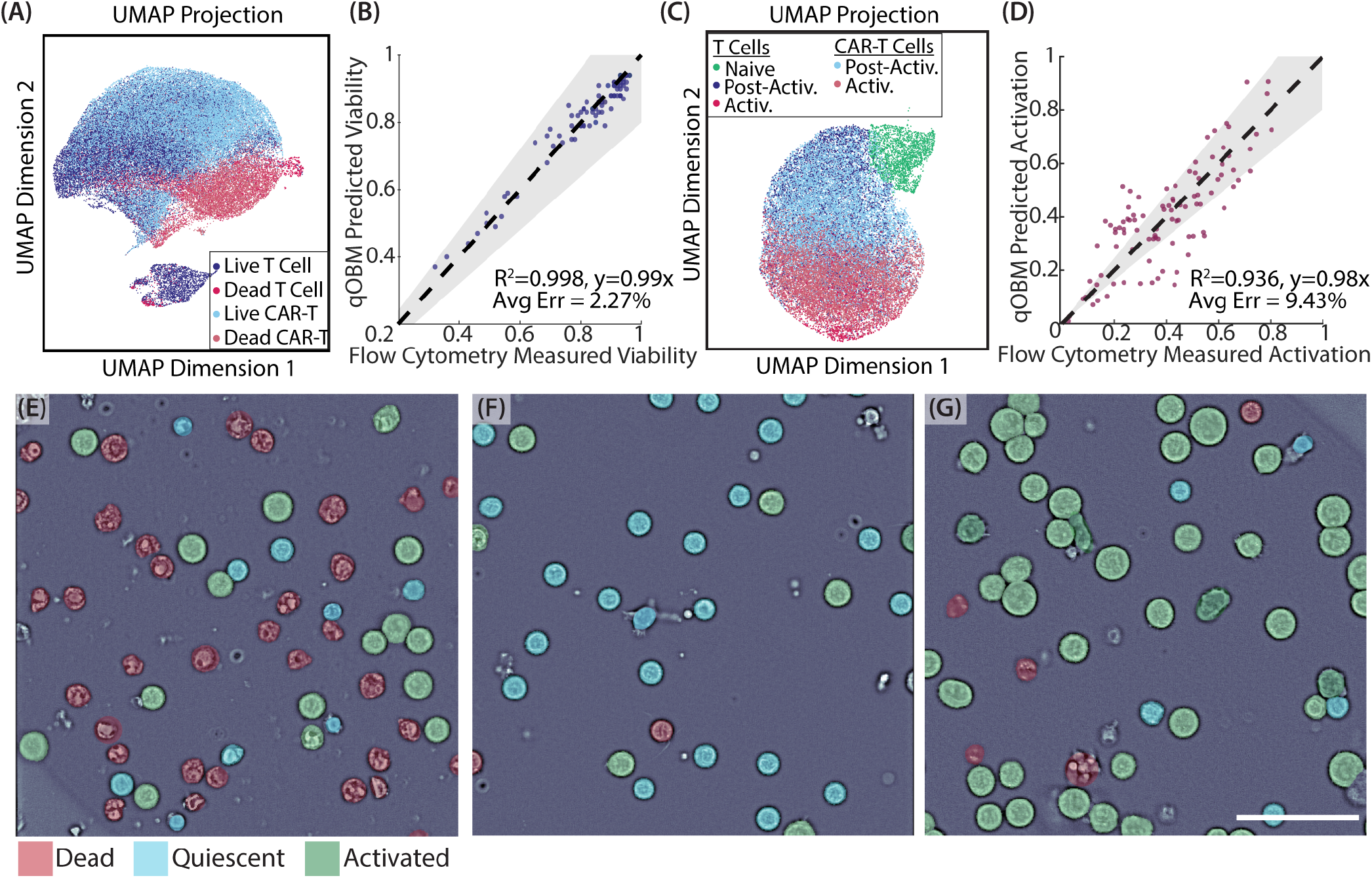
Characterization of the CAR-T cells: (A) UMAP comparing the features from live and dead untargeted T cells to live and dead CAR-T cells. (B) Scatter plot of qOBM predicted viability versus the viability measured with flow cytometry. (C) UMAP comparing the features from activated and unactivated untargeted T cells to activated and quiescent CAR-T cells. (D) qOBM predicted activation versus the CD25^+^ activation measured with flow cytometry. (E)-(J) Images colorized based on cell classification in a cell culture with low viability (E&H), a culture with high viability but low activation (F&I), and a culture with high viability and activation (G&J). Red cells are dead, blue cells are live, quiescent cells, and green cells are live, activated cells. Scale bar is 50 µm.

Next, in further demonstrating the ability of qOBM to characterize CAR-T populations, we applied our activation assay to the live CAR-T cells. Figure 5(C) similarly plots a UMAP of the CAR-T data together with the T cell data which shows a near perfect overlap between the post-activation and activated cells of the two groups (CAR-Ts and T cells). All of the CAR-T cell data included here is from activated or previously activated CAR-T cells; thus, as expected, no CAR-T cells map to the area with naive T cells in green. After running the CAR-T cell data through the qOBM image activation assay (ResNet) (see Figure 5(D)), we again obtain a strong positive correlation with R^2^=0.936 and an average error of 9.43% when comparing the qOBM predicted viability to that of flow cytometry. Based on these results, again from 44 independent donors, we conclude that the activation network can also be applied to the CAR-T cells.

As with the T cells, we can also colorize individual CAR-T cells to visualize the characterization. Figure 5E-G shows cell-by-cell classification in images of cultures with low viability and activation Figure 5(E), high viability but low activation Figure 5(F), and high viability and high activation Figure 5(G). We can see the network outputs in the colorized cells, with red indicating dead cells, blue indicating quiescent live cells, and green indicating activated, live cells. Using this visualization, we show how the network can characterize the CAR-T cells on a cell-by-cell level.

### F. Dynamic qOBM can subtype activated T cells as CD4+ or CD8+ cells with limited accuracy

Finally, we attempted to subtype individual, activated T cells into their function in the immune system. By and large, activated T cells fall into two functions: helper CD4^+^ cells which are responsible for recognizing pathogens and signaling the immune response for the specific pathogen, and killer CD8^+^ cells which are responsible for the immune cytotoxic effects to kill the pathogen.

In an effort to subtype individual activated T cells as CD4^+^ or CD8^+^ cells, we found that phase values alone were not sufficient to discern differences between the functions. A network trained on phase values alone yielded the predictions seen in the confusion matrix in Figure 6(G), which shows a near 50% accuracy for both groups. This indicates the network is effectively guessing at random between CD4^+^ and CD8^+^ cells. Previous work has shown that metabolic differences between CD4^+^ and CD8^+^ cells may be used to characterize activated T cells (18). As such, here we employed dynamic qOBM (DqOBM) to conduct functional imaging of CD4^+^ and CD8^+^ T cell isolations (27, 44). Here, we take a stack of 600 sequential qOBM images taken at 8 Hz (this frame rate was selected to capture metabolic activity within the cell), as seen in Figure 6(A). For each pixel of the image, we can look at the pixel-wise changes in phase (refractive index) and take a Fourier transform of each temporal signal, as seen in Figure 6(B-C). Because the temporal dynamics show a frequency response that seems to follow a power-law decay behavior (see 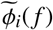 lines seen in Figure 6(C)), we opt to analyze the signals using phasor analysis. In phasor analysis, frequency response of the temporal signal 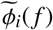 of each pixel is decomposed into two variables termed g and s by obtaining the cosine and sine transform (or the real and imaginary parts of the Fourier Transform). Then the g and s values of each pixel are used to construct a two-dimensional histogram (termed a phasor plot), as seen in Figure 6(E). Further, we can reconstruct a g and s image based on the g and s values (Figure 6F).

**Fig. 6.**
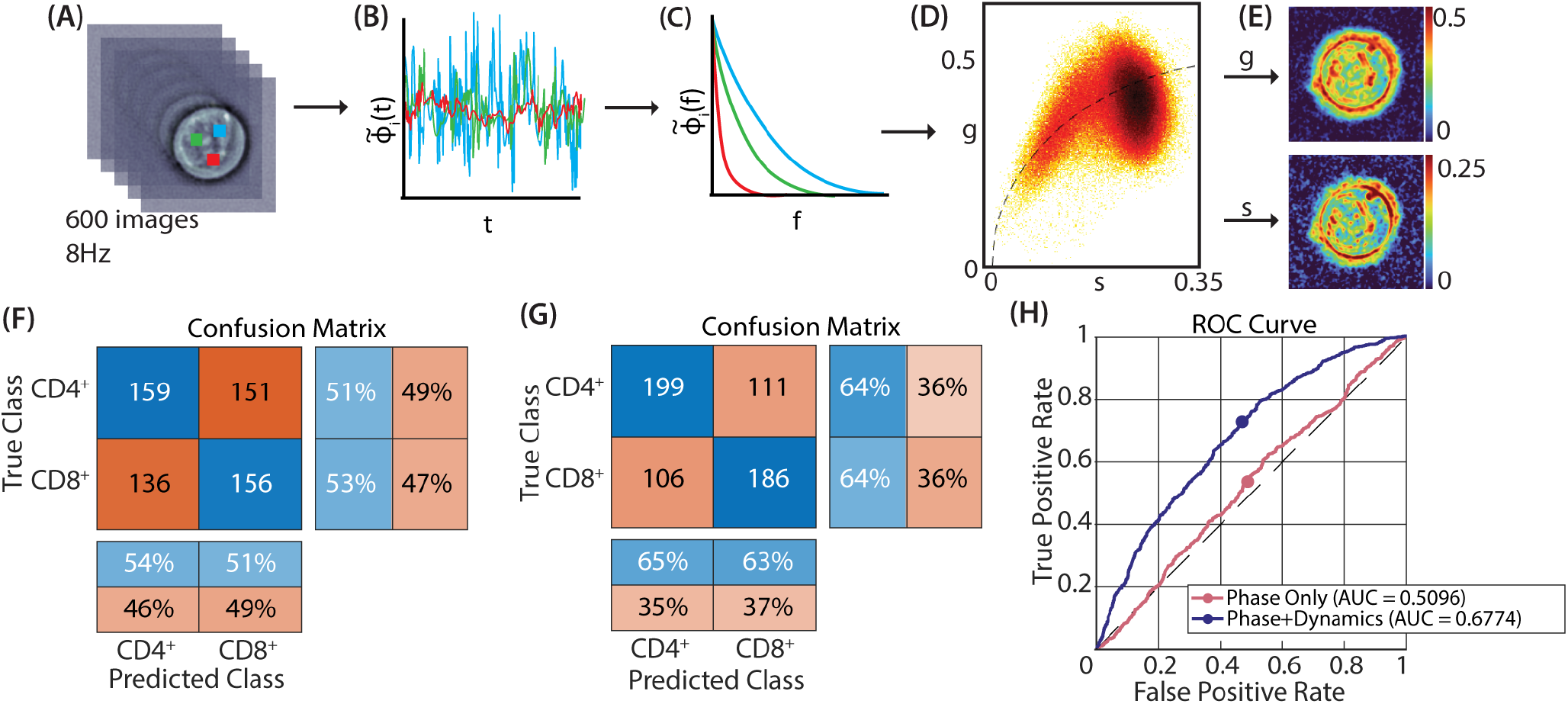
Subtyping of CD4^+^ and CD8^+^ T cells: (A) Stack of 600 quantitative phase qOBM phase images taken at 8Hz. (B) Pixel-wise phase fluctuations from the points indicated by the boxes in (A). (C) The Fourier Transform of the temporal dynamics (lines in (B)) result in decay curves that follow a power-law behavior. (D) 2D histogram phasor plot the frequency response of the temporal dynamics. (E) Functional images based on the g and s values of phasor analysis. (G) Confusion matrix of phase-only classification between CD4^+^ and CD8^+^ cells. (H) Confusion matrix of the phase + dynamics (phasors) classification between CD4^+^ and CD8^+^ cells. (I) ROC curves showing the performance of networks trained only using phase values (red) and trained using phase and dynamic values (blue).

Next, we trained several networks to differentiate between activated CD4^+^ and CD8^+^ cells. To train and test the models we used isolated activated (CD25^+^) Day 3-10 CD4^+^ or CD8^+^ cells with 99+% purity from 3 separate donors. A total of 668 isolated CD25^+^/CD4^+^ and CD25^+^/CD8^+^ cells were imaged from the 3 donors. First, a similar Residual Neural Network to that applied to classify activated cells was trained and tested using the phase, g, and s images as input (see Supplemental Figure S4); however, this network failed to discern any difference between CD4^+^ and CD8^+^ cells. Given the limited data set, we also trained a network based on curated image features. The network was given a total of 146 features from the phase, g, and s images (as described in the methods section). This feature-based network was able to classify CD4^+^ and CD8^+^ cells with 63.99% accuracy (a confusion matrix summarizing these results is seen in Figure 6(H)). Further, we show a receiver operating characteristic (ROC) curve of the model on the testing dataset in Figure 6(I) (additional ROC curves resulting from alternative classification models can be seen in Supplemental Figure S5 & an explanation of the features used to train this model can be seen in Supplemental Figure S8). The resulting area under the curve (AUC) was 0.6774 for the prediction between CD4^+^ and CD8^+^ cells. This ROC plot also compares the performance of this model with cellular dynamic information to one only given phase images of the cells, which shows no ability to differentiate the two activated T cell types (AUC ≈ 0.5). While including dynamic information does demonstrate an improvement over a network trained only on phase values, it is certainly not sufficient for accurate cell subtyping (see Supplemental Figure S9 for a further discussion of the subtyping of CD4^+^ and CD8^+^ cells).

## Discussion and Conclusion

In this work, we demonstrate the ability to monitor T cell cultures in-line in a label-free and non-destructive manner using quantitative phase imaging with qOBM. First, we showed the ability of qOBM to visually identify and quantify cultures growing at high and low rates of cell proliferation, as well as to visualize cell culture contamination from contaminants like yeast and bacteria. Next, we develop residual networks using qOBM images of T cells (i.e., imaging-based assays) to provide accurate estimations of cell culture viability and activation. We show that these networks can be applied in-line to characterize cells on a cell-by-cell basis to estimate culture viability and activation in near real-time. Data were tested using cells from 10 independent donors at-line and 3 in-line with results showing near unity correlations with ground truth flow cytometry measurements. Finally, we also demonstrate that this imaging pipeline can be applied to characterize culture viability and activation of genetically modified therapeutic CAR-T cells. Data were tested using 43 independent donors and the results showed very high agreement with flow cytometry measurements. This process is fully automated, label-free, and can be performed without the need to open the culture environment to perform measurements (i.e., in-line). This marks a significant improvement over end-point analyses like flow cytometry, which are laborious and require that cells be removed from the culture vessel, which can hinder culture expansion and can introduce contaminants. We highlight that the developed in-line qOBM image-based assays can accurately characterize cell cultures relatively quickly with low error when imaging 100+ cells per culture assay. With these conditions, we show errors < 10% compared to flow cytometry, and the image-based assays can be analyzed in < 1 minute to provide near real-time feedback. We expect error can be reduced by imaging more cells per assay, with the trade-off of longer acquisition and processing times.

In this paper, we characterize T cell samples throughout the entire culture process including early and late stage activation, including when the cultures returned to a quiescent state after previously being activated. Other T cell characterization studies have demonstrated the ability to separate naive cells from activated cells but have not included a characterization of cells when they return to dormancy (18). Here we find that there is a phenotypical difference between the two quiescent states (Supplemental Figure S2), and we argue that the post-activation state is of biologic relevance for culturing cell therapeutics as these cells can be introduced in therapeutics as memory cells to train the immune system to respond to disease states and pathogens. Thus, in order to accurately characterize an in-line culture, we recommend that networks be trained on both naive and post-activation quiescent cells to fully capture all states of the T cell cultures.

In this work we used qOBM because it enables quantitative phase imaging using a simple and compact system that is particularly well-suited for in-line cell culture imaging/characterization. Specifically, the incubator-compatible qOBM system is 8in x 10in x 12in, which is extremely compact and suitable for nearly any cell incubator. All microscope components included within the incubator are graded to withstand the humidity and temperatures within the incubator, and we have seen no issues (like condensation on the camera, etc.) that would hinder an in-incubator system. We also note that any existing brightfield microscope (a staple in most cell culture labs) can be adapted to a qOBM system for < $500 (see Supplemental Table S1)(26), making this approach highly accessible. Nevertheless, it is also important to highlight that any QPI embodiment may be implemented to achieve the immune cell characterization described here (some differences may be expected between QPI systems that use coherent illumination vs those, like qOBM, that use partially coherent illumination).

In this work, we also attempted to further sub-type CD4^+^ and CD8^+^ T cells. Based on recent works that utilize the differences in the metabolic activity between CD4^+^ and CD8^+^ cells (18), we hypothesized that functional imaging with DqOBM could also reveal functional differences between the cell subtype. Unfortunately, results showed a very limited ability to differentiate between the two groups. However, parallel efforts using label-free molecular imaging with deep-UV microscopy show that T cell dynamics (similarly analyzed using phasor analysis) clearly differentiate between CD25^+^/CD4^+^ and CD25^+^/CD8^+^ T cells (45). These preliminary in-vitro efforts suggest that the resolution and the coherence properties of the light source (in addition to molecular sensitivity) play a role in the ability to differentiate between the two groups, which implies that coherent QPI alternatives with high-resolution may potentially improve the ability to distinguish CD25^+^/CD4^+^ and CD25^+^/CD8^+^ T cells using phase contrast.

In summary, we have demonstrated a label-free in-line characterization pipeline for immune cell cultures, which can have significant implications for cell manufacturing processes to better enable quality by design. While the analysis pipeline demonstrated in this paper was primarily used to study in-line cultures of T cells and genetically modified CAR-T cells, the approach can also be applied to characterize a wide variety of cell cultures grown in suspension. This low-cost, label-free,and non-invasive in-line imaging pipeline can be broadly applicable to ultimately improve the efficiency of cell manufacturing.

## Methods

### G. Quantitative Oblique Back-Illumination Microscopy

The qOBM setup comprises a conventional brightfield microscope integrated with a modified illumination arrangement, as outlined in previous studies (21, 22, 46, 47). In contrast to the traditional transmission-based illumination utilized in both brightfield microscopy and QPI, the qOBM approach employs an epi-mode illumination strategy using a quartet of LED light sources emitting at 720 nm. These LEDs are strategically positioned around the objective at 90° intervals, as depicted in Figure 1A, and are coupled via optical multimode fibers. In the epi-mode configuration, ∼ 45 mW of light is directed onto the sample. Within the sample, photons undergo multiple scattering interactions, resulting in changes in their trajectory, with a subset being redirected back toward the microscope objective. This phenomenon effectively establishes a virtual light source within the sample, leading to oblique back-illumination, a phenomenon previously termed oblique back-illumination (48). Fluctuations in the refractive index across the sample guide the light either towards or away from the microscope objective. This refractive index-induced modulation induces intensity variations that encode the sample’s refractive index properties. In our investigations, we employ Nikon S Plan Fluor LWD 60X (numerical aperture 0.7) objective.

To achieve quantitative phase imaging with qOBM, we first subtract the intensity images obtained from opposing illumination angles (these intermediate images are called differential phase contrast or DPC images). Subsequently, two orthogonal DPC images, acquired through a total of four distinct acquisitions, are deconvolved using the optical transfer function of the system. This process, previously elaborated upon in references (21–23, 46, 47), ultimately yields quantitative phase contrast images. Leveraging the rich quantitative phase information acquired, we achieve the capability to both visualize and quantify cellular and subcellular structures within the sample, enabling comprehensive tracking of their developmental progression over time.

### H. Compact qOBM System

The compact, incubator-compatible qOBM system operates on the same principles as the system described above. Several substitutions are made to make this system low-cost and suitable for the incubator. First, the pco.edge camera is substituted with a Basler Ace acA3088-57um USB 3.0 Monochrome Camera. The stage here is controlled with a simple XYZ stage (ThorLabs LX30) and stepper motors / motor drivers mounted within the incubator. The LEDs and all electronic controls are mounted outside the incubator. The multimode fibers deliver the LED light to the sample and all electronics are triggered via cables fed through the incubator port. An external PC is used to control all components both within and outside of the incubator.

### I. Dynamic Qobm

To conduct functional imaging with qOBM, we have developed Dynamic qOBM (DqOBM) (49). In DqOBM, a sample is imaged over a period of time – for all samples in this work, this time period is 75 seconds at 8 Hz to obtain 600 images. From these images, we obtain a pixel-wise dynamic frequency response given by the absolute value of the Fourier transform of the temporal phase signal, 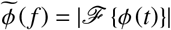 for each spatial pixel in the image. We observe that the frequency response of the pixels exhibits an exponential or multi-exponential decay indicative of subcellular mass movement that more prominently oscillates at low frequencies and dampens exponentially with increasing frequency. This functional behavior is expected for cell structures such as cell membranes (50) and mitochondria (51), among other structures (52). As expected, the dynamic response in background regions shows a mostly flat near-zero amplitude dynamic response, indicative of static behavior. To visualize the cell dynamics, we utilize phasor analysis (53, 54), as it aligns well with the exponential nature of the dynamic phase frequency response, 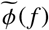.Phasor analysis is a widely used technique for examining spectral and dynamic signals, especially those with exponential characteristics, such as those encountered in fluorescent lifetime and pump-probe microscopy (53, 54). This method involves decomposing signals into two variables, typically referred to as g and s, which are derived from the cosine and sine transforms (real and imaginary components of the Fourier Transform) of the dynamic signals (here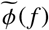) for each spatial pixel in the image at a particular period, τ. In our case, we choose τ=0.5 s depending on the net acquisition rate to decompose the signals into g and s following Eq. 1 and 2, respectively:

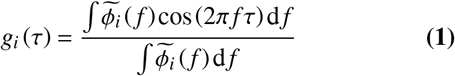

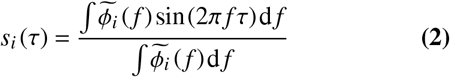

In phasor space, the two components, g and s, act as coordinates, uniquely defining each pixel in the image. As a result, the phasor plot represents a 2D histogram that captures the distribution of g and s values across the image. Similar dynamic signals tend to cluster together in this space, while mixtures of exponential signals create linear mappings between regions.

### J. Isolated Cell Culture

Primary human CD4^+^ and CD8^+^ cells were obtained from an anonymous donor blood (IRB H20288) & commercially available CD3^+^ cell lines (Astarte & Charles River Labs) – Supplemental Tables S2-S4 summarize the donor source. After density gradient centrifugation (Stemcell Technologies 85460) and CD4^+^ (Stem-cell Technologies 17952) and CD8^+^ (Stemcell Technologies 17953) positive isolation. Cells were initially cultured at 0.7e6 cells/ml and maintained at 0.7-2e6 cells/ml in human T cell media consisting of X-VIVO 10 (Lonza 04-380Q), 5% human AB serum (Valley Biomedical HP1022), 10 mM N-acetyl l-cysteine (Sigma A9165) and 55 µM 2-mercaptoethanol (Sigma M3148-100ML) supplemented with 50 units per ml human IL-2 (BRB Preclinical Biologics Repository). Cells were activated 3:1 αCD3/28 Dynabeads (Thermofisher 11161D) to cell ratio on day 0, and beads were removed on day 7. Figures 2, 3, and 4 all include cells from 3 separate donors.

### K. CAR-T cell Culture

CAR-T cells from 43 donors were obtained from a clinical trial (43). Cells were analyzed both freshly thawed and after 24 hours of culture in X Vivo 15 supplemented with 10% FBS, 50 units per ml rh IL-2 and 0.5 ng per ml rhIL-15.

### L. DAPI Staining

Equal parts of cells suspended at a concentration of 1M/ml and 4’,6-diamidino-2-phenylindole (DAPI) stain were combined and well mixed. The mixture was pipetted onto a slide and imaged with 720 nm illumination qOBM imaging. After a qOBM image was taken, a fluorescence image was taken with excitation at 385 nm. A high pass filter was used to exclude any of the excitation light and capture the fluorescent emission. After imaging, fluorescence and phase images were overlaid to label cells as alive or dead.

### M. Bioreactor Cell Culture & Flow Cell Integration

CD3^+^ T cells are seeded into a PBS Mini 0.1L Vertical Wheel (VW) bioreactor with bottom port in 10 mL complete media at a concentration of 1 M cells/mL. These cells are activated for 3 days using TransAct™ (Miltenyi Biotec) in a static environment, then fed up to 70 mL to enable mixing and suspension of cells at 50 rpm. The 70 mL minimum reactor volume submerges the end of the 1/8” ID silicone Masterflex outlet tubing connected to the custom VW bioreactor cap, which enables a peristaltic pump to draw media and cells out into a circulation loop for cell imaging. The cells are pumped out at a flow rate of 8 mL/min through the tubing, sampling valves, and connectors into the custom-designed PDMS flow cell designed for imaging.

After 5 minutes of circulation a representative density of cells has collected in the flow cell and the imaging is completed. During imaging, both inlet and outlet tubing to the flow cell are clamped and the incubator turned off to prevent vibrations from the pumps and other systems from interfering with cell settling and image quality. After imaging, the cells in the circulation loop are flushed back into the VW via the bottom port by switching a valve and pushing through a slug of sterile air prepared in a sterile syringe, then followed by filtered cell-free culture media using the hollowfiber as a novel active cell filter. Separate input and output locations in the VW ensures that cells circulated for imaging are continually mixed and distinct. A full detailed illustration of the flow cell can be seen in Supplemental Figure S10.

### N. Ground Truth Flow Cytometry Procedure

To stain for activation markers, T cells were stained with CD8 (SK1; Biolegend, RRID: 344722), CD4 (RPA-T4; Biolegend, RRID: 300538), CD25 (M-A251; Biolegend, RRID: 356104), and CD69 (FN50, BioLegend, RRID: 310930) at 1:200 and aqua Live/Dead (ThermoFisher L34966) at 1:500. On the day of staining, cells were washed 3x times with FACS (PBS, 0.1% BSA, 2mM EDTA, pH 7.4), stained with the aforementioned stains for 30 minutes at 4°C, and washed 3x times with FACS. Samples were run on Cytek Northern Lights. Flow cytometry procedures are identical between isolated cell cultures and those grown in the bioreactor.

### O. Network Creation and Testing

For each trained network, 35µm x 35µm images with a binary class (i.e. alive/dead or activated/quiescent) were fed into a residual neural network (ResNet). These images contained the phase values of a single cell centered within the image. Cells were segmented and NaN padded outside of the cell. 70% of the data was used for training while 30% of the data was saved for testing. After the network was trained and tested, additional testing was done as the network was applied to other patient cell lines and other growth conditions. To compare measurements with flow-cytometry measurements, we only utilize cultures with >100 individual cells in order to have enough cells to characterize a sample.

### P. Feature Analysis

Quantitative image features extracted from the qOBM images were analyzed to assess structural differences between different T cell subtypes. To accomplish this, images of individual cells were cropped to fit in the center of a 35µm x 35µm image. For each cell, features were extracted based on texture analysis (55), fractal analysis (56, 57), Fourier space features (58), and mathematical auto-correlation transformations (59, 60). Detailed explanations regarding the computed features can also be found in Ref. (23). Feature selection ranking was performed using Minimum Redundancy and Maximum Relevance, Neighborhood Component Analysis, and the Chi-square tests, as implemented by Matlab’s functions fscmrmr, fscnca, and fscchi2, respectively. Features selected for the correlations with flow cytometry were deemed those with the highest correlations with the flow cytometry values. Figures demonstrating the feature ranking for each network can be seen in Supplemental Figures S6-S8. A figure showing the uniformity of features between donors can be seen in Supplemental Figure S1. In the figures above, we note specific Thresholds to obtain fractal dimension values. Here, Threshold 2 contains refractive indices between 1.6289-1.6292, Threshold 6 contains refractive indices between 1.6294-1.6230, Threshold 7 contains refractive indices between 1.6297-1.6300, and Threshold 14 contains refractive indices between 1.6305-1.6308.

For the CD4/CD8 cell isolations, three-channel images were constructed with the red channel containing refractive index information, the green channel containing “g” dynamic information, and the blue channel containing “s” dynamic information. The same features were extracted for all 3 individual channels, and the same feature selection process was implemented.

## ACKNOWLEDGEMENTS

**Acknowledgments:** Figure 1A was created with https://biorender.com/.

## Funding

We acknowledge the following funding sources:

National Science Foundation Graduate Research Fellowship: NSF GRFP DGE-2039655 (C.E.S. & A.D.S.T.)

Burroughs Wellcome Fund: CASI BWF 1014540 (F.E.R.)

National Institute of General Medical Sciences: R35GM147437 (F.E.R.)

National Institutes of Health Cell and Tissue Engineering Training Program: T32GM145735 (A.D.S.T.)

National Institute of Allergy and Infectious Diseases grant: R01AI171892 (G.A.K) The Billie and Bernie Marcus Foundation: (F.E.R.)

Georgia Tech Research Institute Independent Research and Development (IRAD) funds: (F.E.R.)

Georgia Institute of Technology: (F.E.R.)

## Author contributions

C.E.S., V.G., P.C.C., and F.E.R. designed the experiments outlined in the paper. C.E.S. and P.C.C. acquired the data. C.E.S. designed the flow cell, analyzed the data, and made all figures in the manuscript.

C.E.S. and F.E.R. wrote the paper. V.G. developed the framework for the residual neural network to classify cells. B.K., B.Wang, and B.Wicker. implemented in the in-line T cell culture. A.S. and I.L. isolated and analyzed the quiescent and activated CD4^+^/CD8^+^ populations. R.Q.C. and B.J. developed the masking algorithm for the qOBM imaged cells. L.K. cultured the CAR-T cells used. N.L. ran flow cytometry on the CAR-T cells. C.E.B. provided the CAR-T cells. All authors reviewed the results and approved the final version of the manuscript.

## Competing interests

G.A.K. reports equity or consulting roles for Sunbird Bio, Port Therapeutics, Send Biotherapeutics, and Ridge Biotechnologies. All other authors declare they have no competing interests.

## Data and materials availability

Data and code underlying the results presented in this paper are not publicly available at this time but may be obtained from the corresponding author upon reasonable request. Similarly, materials used in the study are largely outlined in the Methods section; however, further details may be obtained from the corresponding author upon reasonable request.

## Ethics approval and consent to participate

All experimental protocols utilizing human cells were conducted under the guidelines of the Georgia Institute of Technology.

