## Supplemental Text for "Label-Free In-Line Characterization of Immune Cell Culture using Quantitative Phase Imaging"

### Q. Supplementary Text.

#### Cost of the qOBM Microscope.

**Table S1. Detailed outline of the parts necessary to transform an existing brightfield microscope into a qOBM system.** This price is <$500. We also highlight the key components used in this microscope in this study.

| ***Detailed costs to convert a brightfield microscope to a qOBM system*** | | |
|---|---|---|
| **Item** | **Quantity** | **Cost / Unit** |
| Arduino Nano Every | 1 | $13.00 |
| LuxeonStar Far Red Rebel LED | 4 | $8.00 |
| LuxeonStar High Alpha Heat Sink | 4 | $7.07 |
| LuxeonStar BuckPuck DC Driver | 4 | $10.99 |
| Thorlabs Aspheric Condenser Lens, f=16mm, N.A.=0.79 | 8 | $20.56 |
| Thorlabs 0.50 N.A. Step-Index Multimode Fibers (1 meter) | 4 | $31.77 |
| Thorlabs SMA905 Multimode Connector | 4 | $12.46 |
| | **TOTAL** | **$ 458.64** |
| ***Key components of the brightfield microscope used in this study*** | | |
| Basler Ace acA3088-57um USB 3.0 Monochrome Camera | 1 | $419 |
| Nikon Plan Fluor ELWD, 60X, 0.7 N.A. | 1 | $4,975 |

.

DRAFT

### Summary of Donor Numbers.

**Table S2. Detailed enumeration of the number of cells used for training for each network.** This table highlights the number of cells used and the number of donors these cells came from.

| *Number of cells contained in each training set for network formation* | | |
|---|---|---|
| **Group** | **Number of Cells** | **Number of Donors** |
| Alive | 462 | 2 |
| Dead | 72 | 2 |
| Activated | 1,937 | 3 |
| Quiescent | 952 | 3 |
| $CD4^+$ | 795 | 3 |
| $CD8^+$ | 762 | 3 |
| Naive | 732 | 3 |
| Post-Activation | 533 | 3 |

**Table S3. Detailed enumeration of the number of cells used for testing for each network.** This table highlights the number of cells used and the number of donors these cells came from.

| *Number of cells contained in each testing set for network formation* | | |
|---|---|---|
| **Group** | **Number of Cells** | **Number of Donors** |
| Alive | 33,443 | 10 |
| Dead | 5,474 | 10 |
| Activated | 7,501 | 11 |
| Quiescent | 11,251 | 11 |
| $CD4^+$ | 341 | 3 |
| $CD8^+$ | 327 | 3 |
| Naive | 314 | 3 |
| Post-Activation | 10,937 | 11 |

**Table S4. Detailed enumeration of the number of cells used from each donor & the source of the donor cells.** This table highlights the number of cells used from each donor, the source of the cells, and the ages of the cell culture imaged.

| *Number of cells imaged per each donor* | | | |
|---|---|---|---|
| **Donor #** | **Cell Source** | **# Cells** | **Culture Ages Imaged (Days)** |
| 1 | Astarte Human $CD3^+$ T cells | 12,827 | 5-14 |
| 2 | Charles River Labs $CD3^+$ T cells | 2,879 | 0-14 |
| 3 | Charles River Labs $CD3^+$ T cells | 14,250 | 0-10 |
| 4 | Healthy Donor Peripheral Blood Draw Isolation | 1,837 | 0-21 |
| 5 | Healthy Donor Peripheral Blood Draw Isolation | 1,400 | 0-21 |
| 6 | Healthy Donor Peripheral Blood Draw Isolation | 1,147 | 0-21 |
| 7 | Charles River Labs $CD3^+$ T cells | 489 | 3-10 |
| 8 | Charles River Labs $CD3^+$ T cells | 672 | 10 |
| 9 | Charles River Labs $CD3^+$ T cells | 745 | 3-10 |
| 10 | Charles River Labs $CD3^+$ T cells | 663 | 10-17 |
| 11 | Charles River Labs $CD3^+$ T cells | 992 | 10-22 |

.

DRAFT

### Homogeneity of Cell Culture Features.

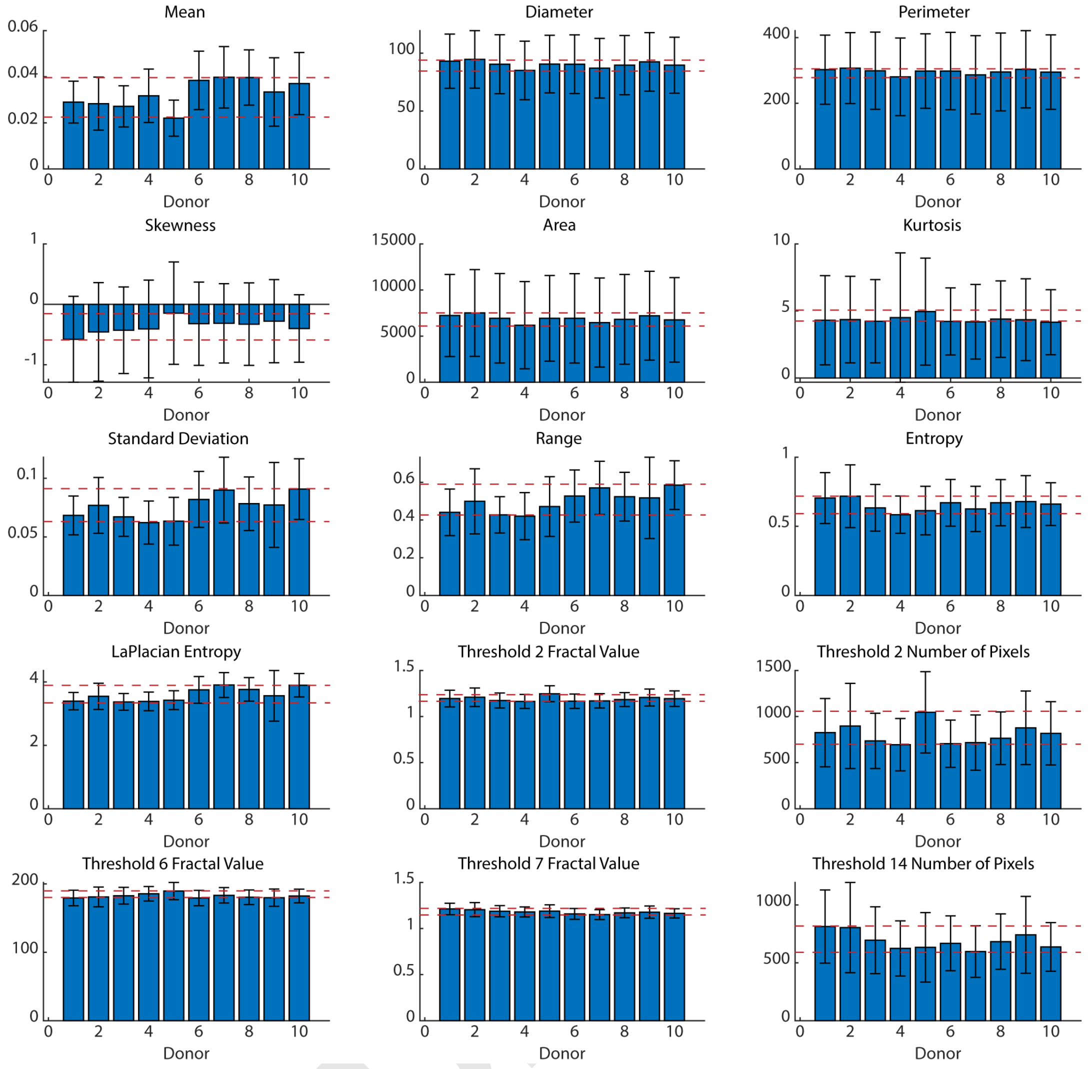


**Fig. S1. Plots here demonstrate the values of features extracted in Figure 2.** Here, we see no major differences between donor cell lines. Nearly all donor lines have a mean within the mean +/- the standard deviation of the populations. The highest and lowest mean of the features are indicated by the red dotted lines. Note that nearly all of the values have a standard deviation that crossed both lines indicating that the value likely falls within this range. Populations here include all cells (alive and dead). Variation here may arise from fluctuations in culture viability and activation; however, we note that the average viability across a donor (as seen here) in >85%.

.

**Machine learning network to distinguish between quiescent, unactivated T cells and previously activated T cells.**

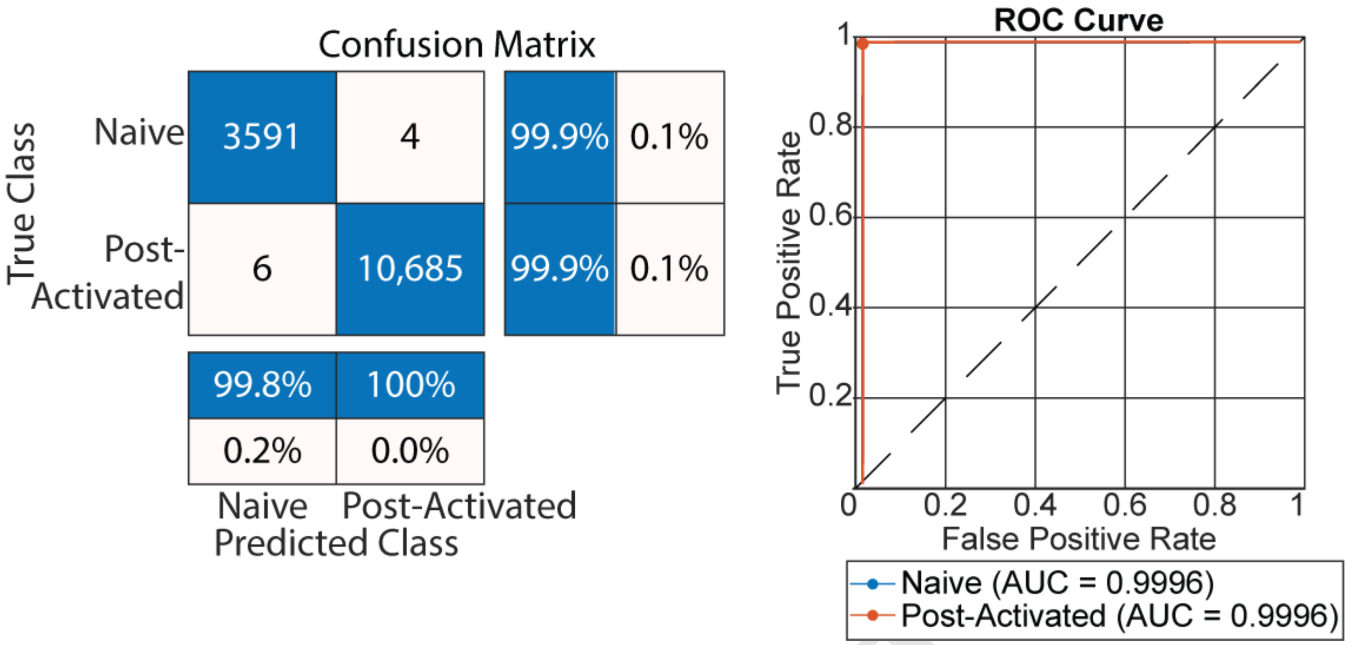


**Fig. S2. Output of a ResNet trained to distinguish between quiescent D0 cells and D21 cells that previously had a culture activation >99%.** We see excellent separation between the D0 and D21 cells with near perfect divisions and an ROC curve near 1.

.

DRAFT

**Statistical analysis of the culture composition.**

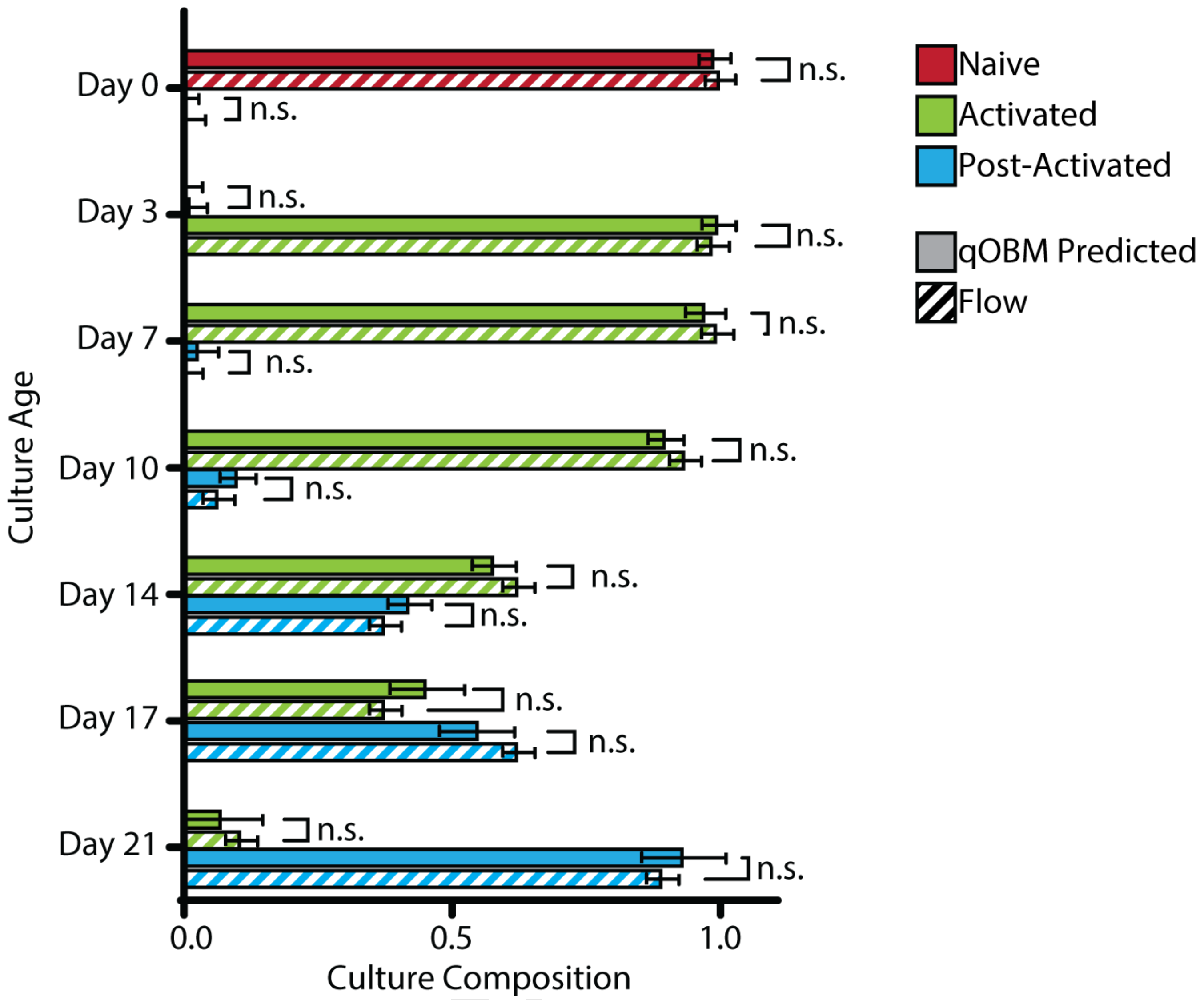


**Fig. S3. Statistical analysis comparing the qOBM predicted composition of a cell culture on a given day with that of flow cytometry.** Here, we see no statistically significant differences between the groups.

.

**Image-Based Residual Neural Network to Differentiate CD4+ and CD8+ cells.**

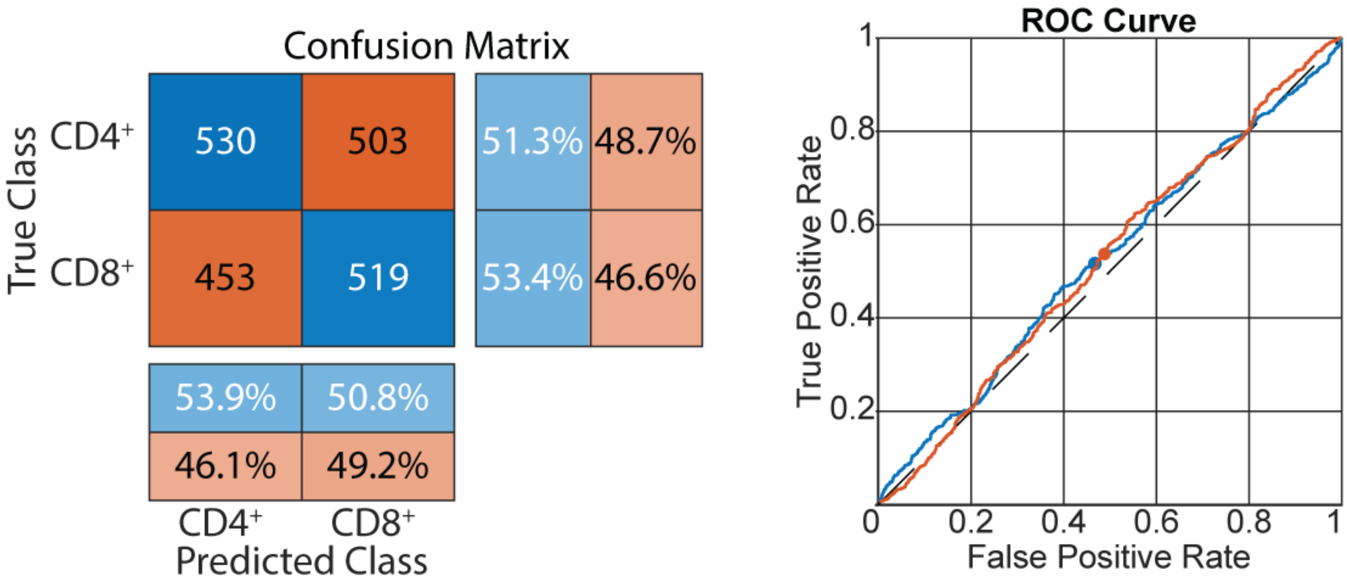


**Fig. S4. Output of a ResNet trained to distinguish CD25+/CD4+ and CD25+/CD8+.** This is done with a purity of >99%. The input of this network was the 3 channel RGB images of cells where the red (R) channel was cell phase image, the green (G) channel was the DqOBM G image, and the blue (B) channel was the DqOBM S image. The residual neural network was not able to differentiate between the populations

.

DRAFT

**ROC Curves for Trained Networks.**

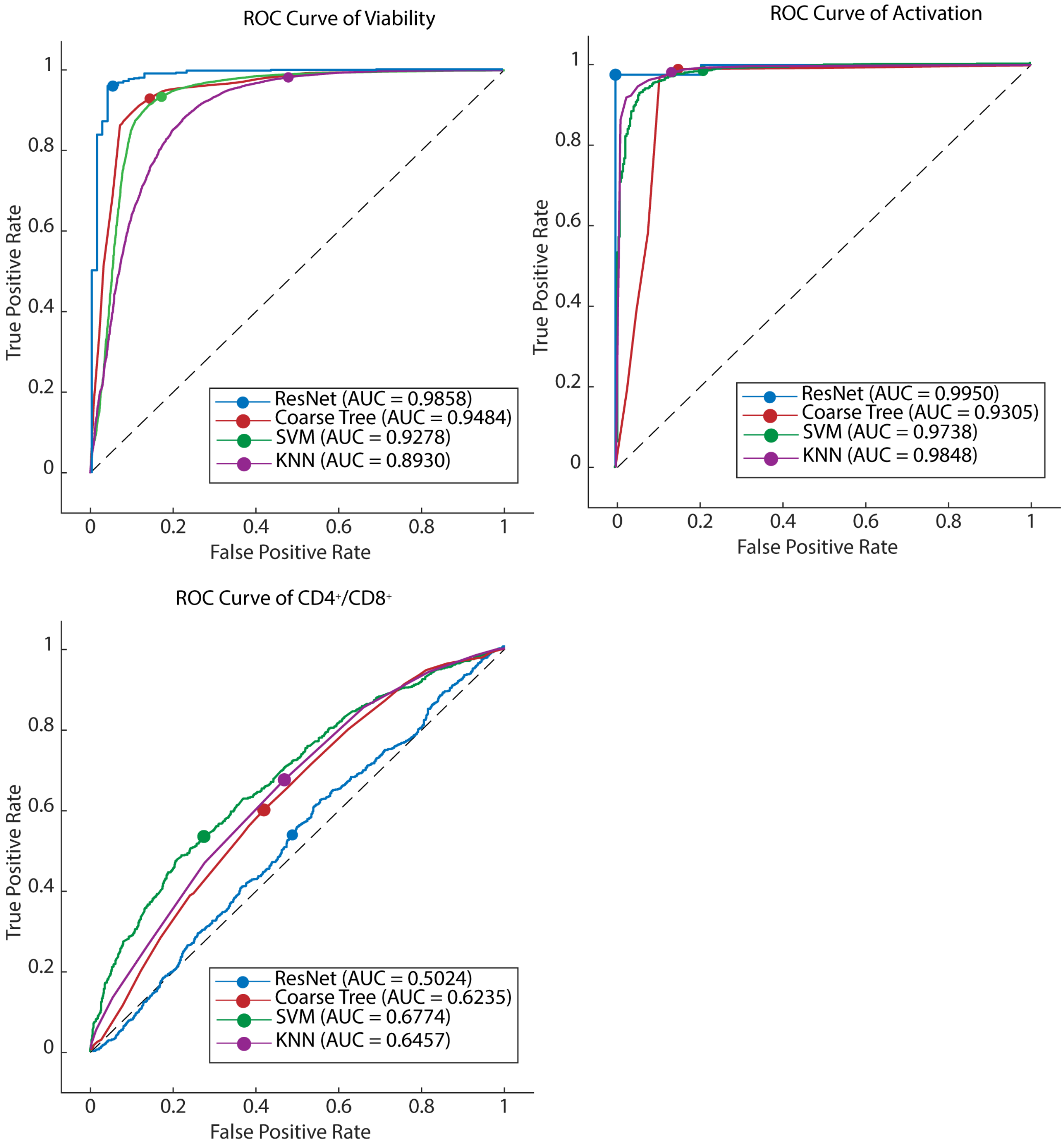


**Fig. S5. Receiver operating characteristic (ROC) curves of the output of different tested networks.** Here we see that the ResNet performs best for viability and activation but can not distinguish $CD4^+/CD8^+$ cells. Here, feature based networks perform better with an SVM performing the best.

.

**Feature Selection.**

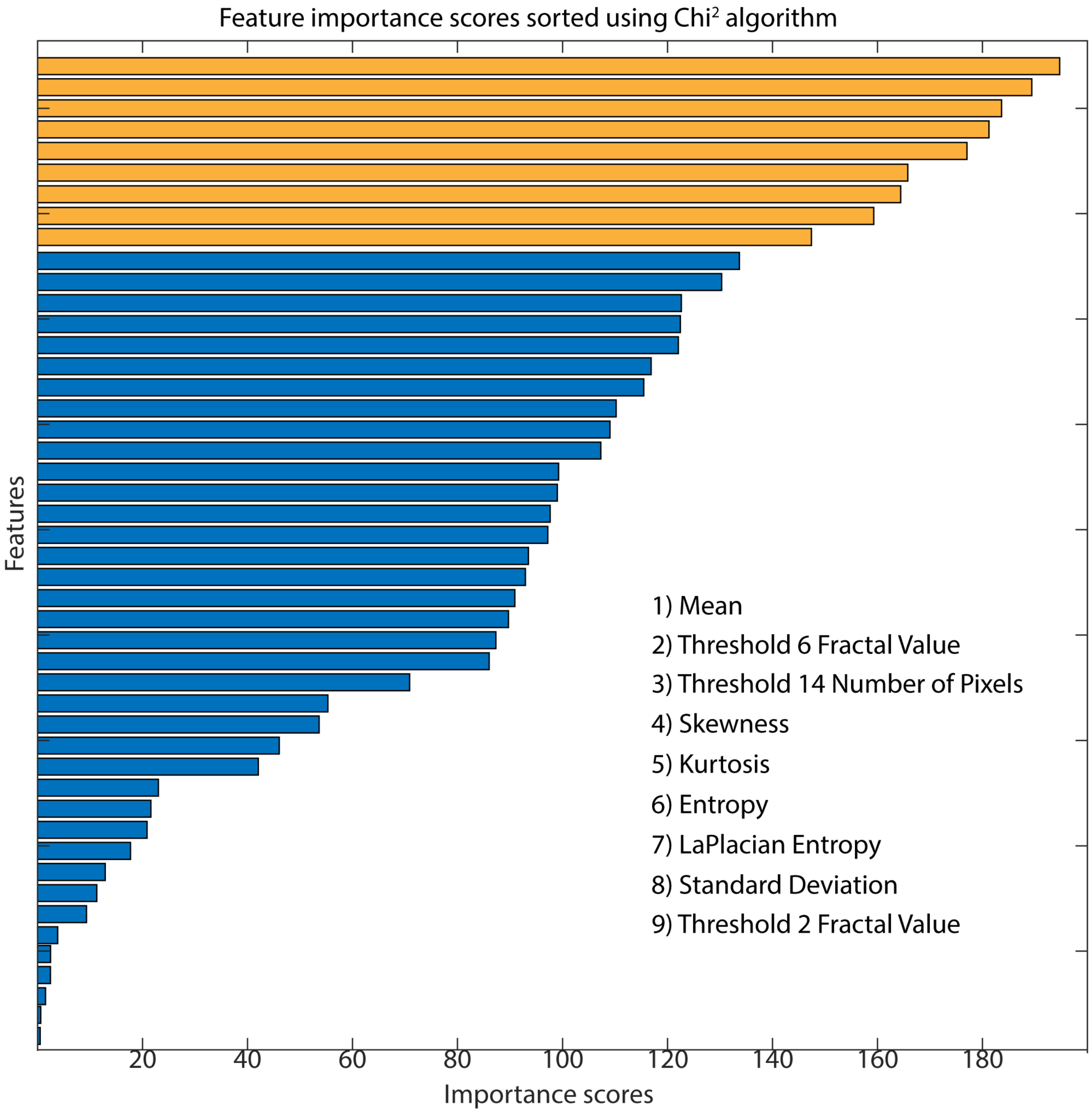


**Fig. S6. Chi$^2$ feature ranking for the viability network.** Here, we can see the Chi$^2$ values for the top 9 features used in the network (highlighted in yellow). Redundant features were removed.

.

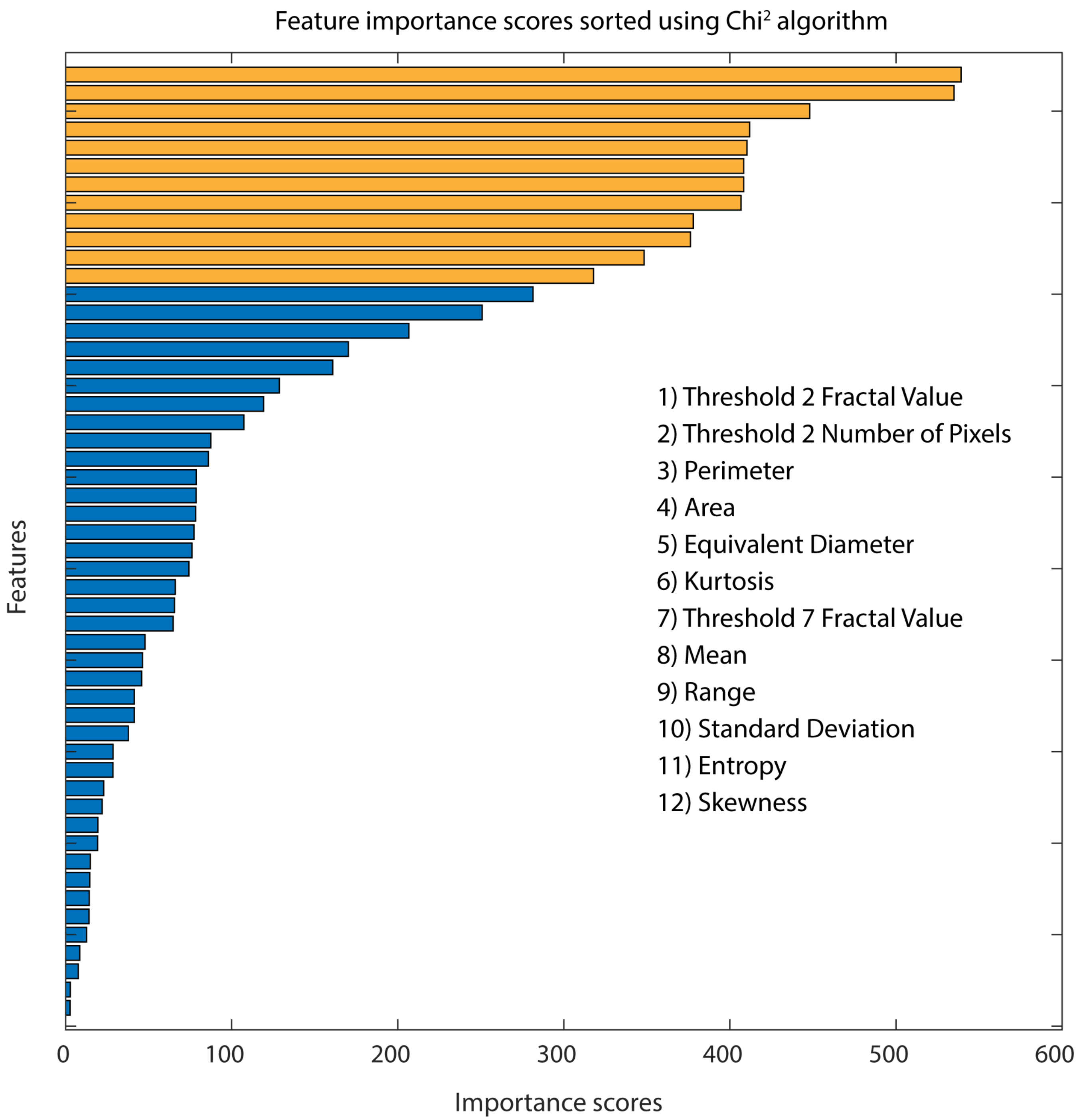


**Fig. S7. Chi$^2$ feature ranking for the activation network.** Here, we can see the Chi$^2$ values for the top 12 features used in the network (highlighted in yellow). Redundant features were removed.

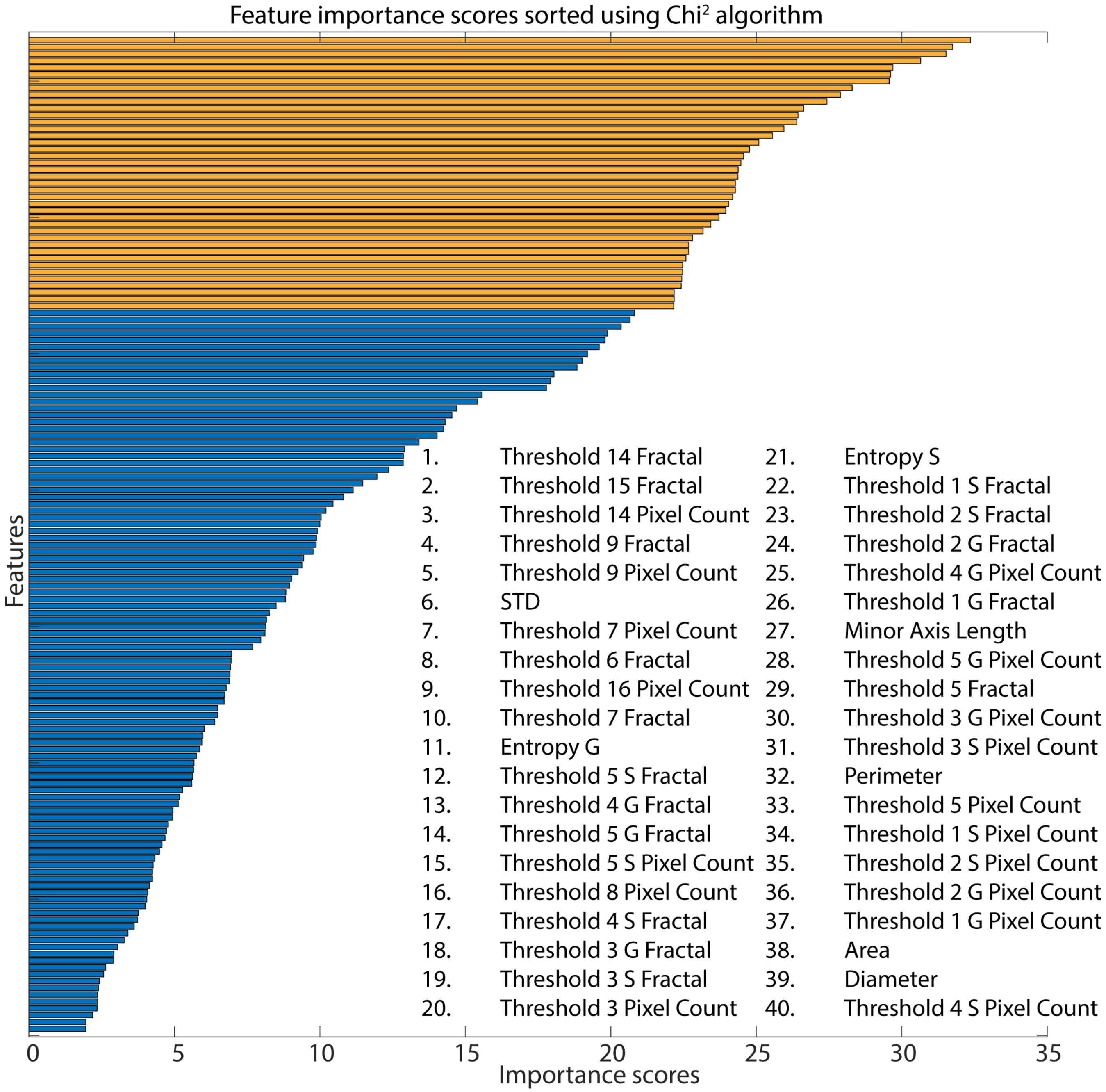


**Fig. S8. Chi$^2$ feature ranking for the CD4$^+$/CD8$^+$ network.** Here, we can see the Chi$^2$ values for the top 40 features used in the network (highlighted in yellow). Redundant features were removed.

#### Residual Neural Network to Differentiate CD4⁺ and CD8⁺ cells.

. To further understand the inaccuracy included by the model, we compared the phasor plots from $CD4^+$ and $CD8^+$ cells (as seen in Figure S9(A)). We see nearly identical high dynamics activity (located farthest away from 0). Perhaps some differences exist in lower regions of dynamic activity; however, much of this is likely due to noise and no true clusters can clearly be distinguished.

Finally, we attempted to apply this model to different donor culture cells from outside of the training and testing data. Here, we do not see any clear correlation between the qOBM predicted $CD4^+$ / $CD8^+$ values and the ground truth flow cytometry values (as illustrated in Figure S9(B-C)). Both exhibit high errors with an average error of 24.01% and 23.53% and $R^2$ values of 0.069 and 0.075 for the $CD4^+$ and $CD8^+$ cultures, respectively. With this, we conclude that the trained qOBM models are currently unable to differentiate between $CD4^+$ and $CD8^+$ cells.

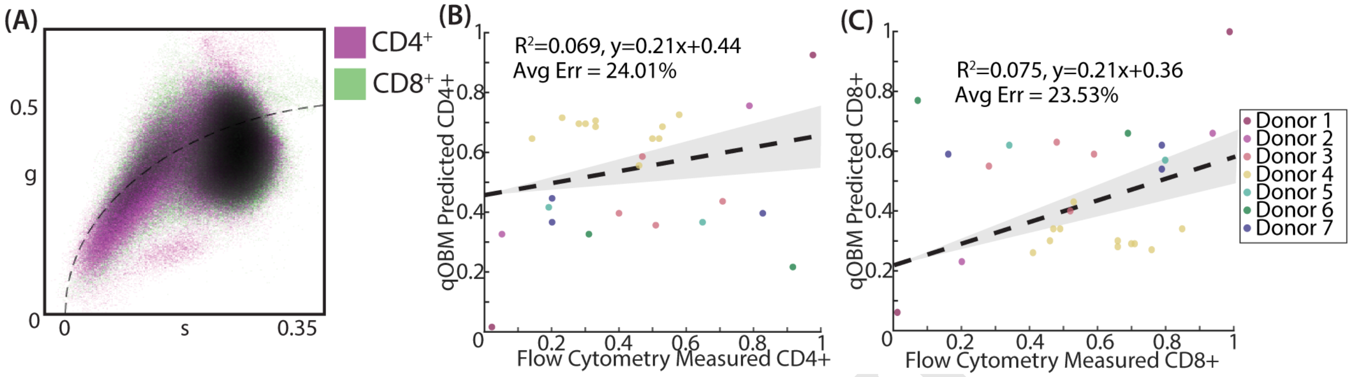


**Fig. S9. Further analysis of phase-mediated $CD4^+$/$CD8^+$ separation in phasor space.** (A) overlaid phasor plots for $CD4^+$ and $CD8^+$ cells showing little distinguishable dynamic active. (B-C) the performance of the network extended to $CD4^+$ cells (B) and $CD8^+$ cells (C).

DRAFT

**Flow Cell Setup & Dimensions.**

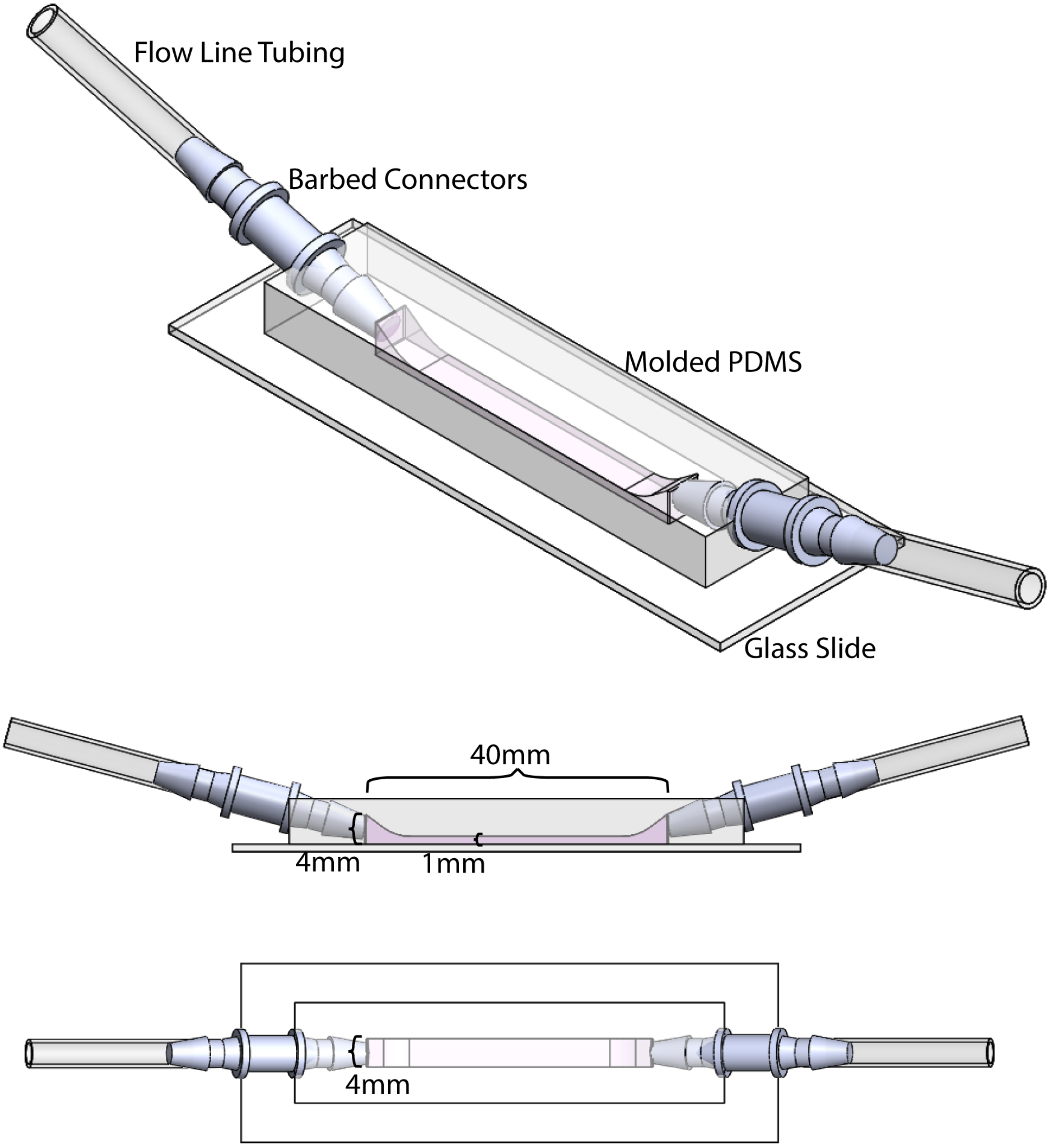


**Fig. S10. Detailed diagram of the flow cell dimensions and setup** Appropriate dimensions at listed in the diagram.

.
